# Confinement, jamming, and adhesion in cancer cells dissociating from a collectively invading strand

**DOI:** 10.1101/2024.06.28.601053

**Authors:** Wei Wang, Robert A. Law, Emiliano Perez Ipiña, Konstantinos Konstantopoulos, Brian A. Camley

## Abstract

When cells in a primary tumor work together to invade into nearby tissue, this can lead to cell dissociations—cancer cells breaking off from the invading front—leading to metastasis. What controls the dissociation of cells, and whether they break off singly or in small groups? Can this be determined by cell-cell adhesion or chemotactic cues given to cells? We develop a physical model for this question, based on experiments that mimic aspects of cancer cell invasion using microfluidic devices with microchannels of different widths. Experimentally, most dissociation events (“ruptures”) involve single cells breaking off, but we observe some ruptures of large groups (*∼* 20 cells) in wider channels. The rupture probability is nearly independent of channel width. We recapitulate the experimental results with a phase field cell motility model by introducing three different cell states (follower, guided, and high-motility metabolically active leader cells) based on their spatial position. These leader cells may explain why single-cell rupture is the universal most probable outcome. Our simulation results show that cell-channel adhesion is necessary for cells in narrow channels to invade, and strong cell-cell adhesion leads to fewer but larger ruptures. Chemotaxis also influences the rupture behavior: Strong chemotaxis strength leads to larger and faster ruptures. Finally, we study the relationship between biological jamming transitions and cell dissociations. Our results suggest unjamming is necessary but not sufficient to create ruptures.

## I. INTRODUCTION

Collective cell migration shows up in many developmental processes, like wound healing [1], embryogenesis [2, 3], and tumor growth and invasion [4, 5]. Compared to single-cell migration, collective migration is often more persistent since cells are more coordinated due to their mutual communications and interactions [5, 6]. The ability of cells to rearrange fluidly in collective migration may be controlled by an unjamming transition [7–10] shown in dense biological tissues, which is similar to the well-known solid-liquid phase transition of materials [7]. This jamming-unjamming transition has been extensively studied using different theoretical models [7–11], which predict that cell tissues can be fluidized by increasing cell motility and deformability [7, 8, 10].

Collective cell migration *in vivo* often occurs within confined spaces [12–14]. For example, during cancer invasion, the primary tumor extends a strand of cells into surrounding tissues, resulting in cells being confined, followed by the detachment of a single cell or a group of cells from the strand. Cancer metastasis is a complex process that requires cancer cells first to detach from the primary tumor either individually or as small clusters, and then migrate through adjacent tissue, circulate in the vasculature, and finally survive and proliferate in distant organs [5]. Though clusters of tumor cells circulating in the bloodstream (usually 2–50 cells) are rare, they can drastically increase metastatic potential compared to single-cell seeding [5, 6]. Moreover, it has been reported that circulating tumor cells may even cluster with white blood cells or platelets to escape immune surveillance [15]. Therefore, there is an urgent need to understand the formation of these tumor clusters.

In this work, we study how single cells and clusters of cells dissociate from an invading front of a tumor under different degrees of geometric confinement. We use the approach we developed in [16] to study invasion and dissociation *in vitro*, with cells induced to invade into a narrow microchannel by a gradient of serum (Fig. 1). In these experiments, we see one or several cells as a cluster detach from an invading stream in a microchannel of controllable width [16]. We have observed that the majority of dissociation events (“ruptures”) are single-cell ruptures, but in wide channels rupture of larger clusters is more common. Interestingly, the timescale of rupture appears to be independent of channel width in the experiments. We construct a model of this dissociation process by describing the cells as deformable crawling objects using a phase field model [7, 17]. This model allows us to describe the shape of each individual cell, which is key to unjamming [7–10], while allowing us to model cells detaching from one another. By incorporating different cell states and cell-wall adhesion into the standard phase field model, we find that our model recapitulates the cluster-size distribution and survival probability observed in experiments, but this requires the presence of leader cells and a decrease in intercellular adhesion under confinement. Subsequently, we investigate the impact of four critical parameters in our model—cell-cell adhesion strength, cell-wall adhesion strength, chemotaxis strength, and the number of leader cells—on the rupture behavior of invading monolayers in confinement. In particular, we find that increasing cell-cell adhesion makes dissociation slower, but also increases the average size of the dissociating cluster. Leader cell formation probability controls the balance between single-cell and multiple-cell dissociations. Furthermore, we explore the relationship between cell dissociation behavior and the unjamming transition, finding that unjamming is necessary but not sufficient to create ruptures.

**FIG. 1.**
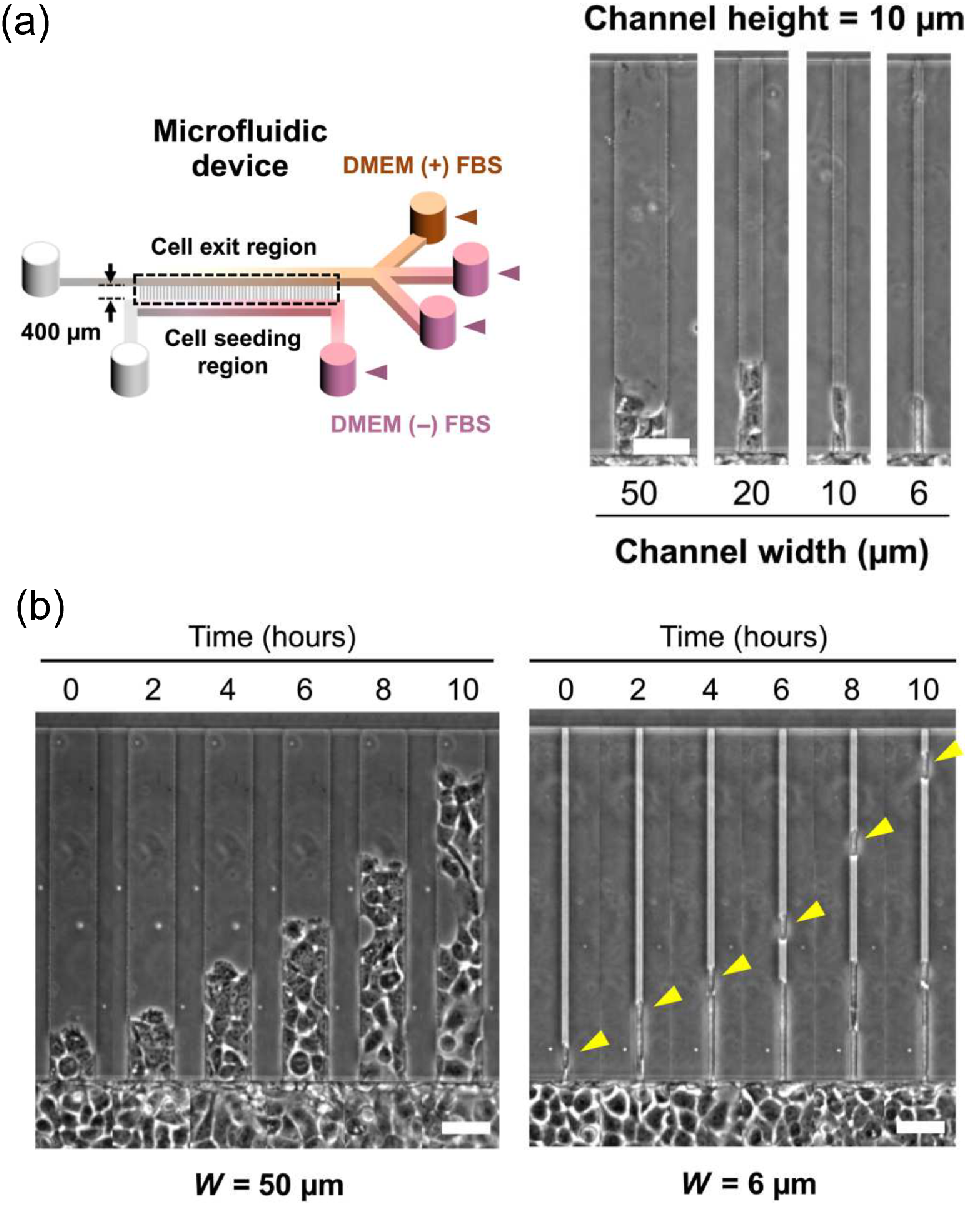
Experimental setup. Reprinted from Ref. [16]. (a) Schematic of microfluidic device with microchannels of prescribed dimensions and phase-contrast images of microchannels of different widths. Cells were cultured in Dulbecco’s modified Eagle’s medium (DMEM). (b) Representative phase-contrast snapshots of invading human A431 epidermoid carcinoma cells confined in 50 *μ*m and 6 *μ*m microchannels. Yellow arrowheads indicate an ongoing single-cell dissociation. Cells are exposed to a gradient of fetal bovine serum (FBS); FBS concentration increases in the +*y* direction. Scale bars, 50 *μ*m.

## II. EXPERIMENTAL RESULTS

We begin with a focus on our recent experiments using Human A431 epidermoid carcinoma cells in microchannels [16], where confluent cell monolayers follow a gradient of chemoattractant (fetal bovine serum, FBS) to enter the microchannels. These experiments revealed details of the biochemical mechanism of cell dissociation from collectively migrating strands in confinement [16]. Figure 1(a) shows experimental snapshots for different microchannel widths. We can observe that cells are entering microchannels from the seeding region (the wide region with cells at the bottom) and following the chemical gradient. We see that a single cell or clusters of several cells dissociate from the main bulk of the cell collective; we show an example of a single-cell dissociation in a 6-micron channel and a dissociation of a large cluster in Fig. 1(b).

We re-analyze experiments of invasion in different channel widths first presented in Ref. [16], measuring the sizes of clusters breaking off and the times of cluster ruptures. We plot the distributions of cluster size as a function of microchannel width in Fig. 2(a). The mean cluster size *A*, plotted as dashed lines in Fig. 2(a), increases with channel width *W*. We see that across all the channel widths, though cluster size increases, the most common outcome is a single-cell dissociation. The probability of having a *n*-cell rupture decreases sharply with increasing *n*. However, in wide channels (20 *μ*m and 50 *μ*m), we do see a good number of larger ruptures [Fig. 2(a)]. In fact, in the 50-micron channel, observing clusters of *>*10 cells is the second most probable category. For full histograms without categorization, see Appendix F.

**FIG. 2.**
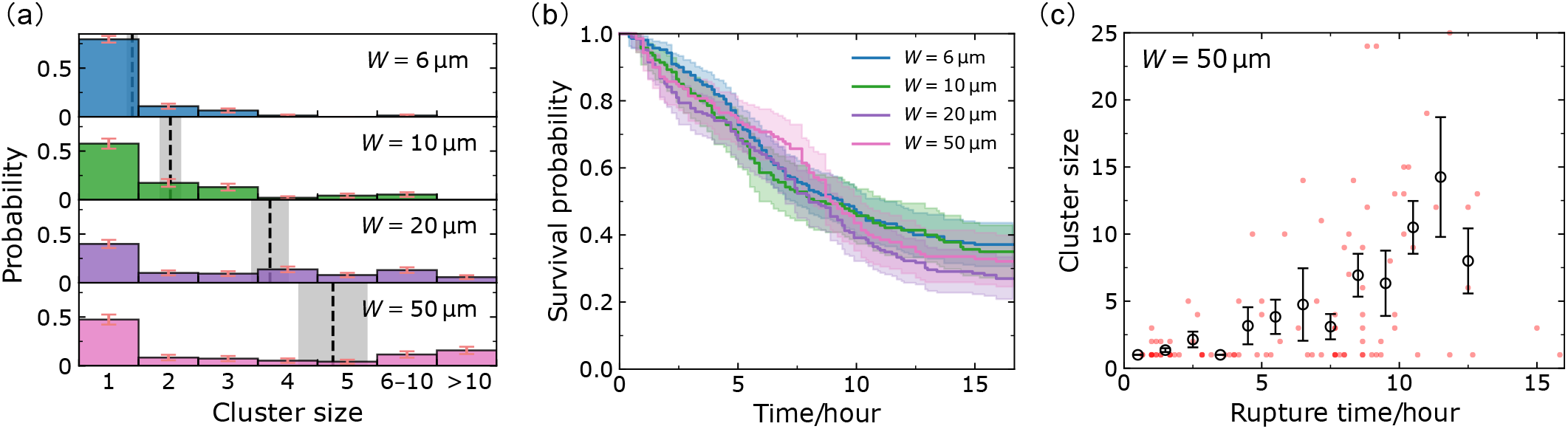
Analysis of experimental dissociation events. (a) Dissociation cluster size distributions for *W* = 6, 10, 20, and 50 *μ*m microchannels. The dashed lines and gray areas denote the average cluster sizes *A* = 1.4, 2.0, 3.7, 4.8, and the associated standard errors of the mean (mean *±* SE). Note that we are using cluster size categories 6–10 and *>*10 to group together similar-sized large clusters; full histograms are shown in Fig. S1(a). Panel (b) shows the survival curves for different channel widths extracted by Kaplan-Meier survival analysis [18]. The shaded areas represent the 95% confidence intervals; the final survival probabilities are *K*_*s*_ = 73*/*210 (35%), 48*/*140 (34%), 51*/*189 (27%), 45*/*140 (32%), respectively; the denominators of *K*_*s*_ represent the number of assays in the experiment. (c) Scatter plot in the (rupture time, cluster size) phase plane. Data points (red) are plotted with transparency; darker points indicate ruptures occurring in more than one assay with the same cluster size and rupture time. The circles represent the average cluster sizes within each time bin (1 hour), and the error bars indicate the corresponding standard errors. Figures for all channel widths are given in supplementary Fig. S1(b).

How quickly do these ruptures happen? We characterize the timescale for the first rupture to happen by measuring the survival probability *S*(*t*), defined as the probability that the monolayer in a microchannel has not ruptured at time *t*. By comparing *S*(*t*) curves for different channel widths, we can see how fast ruptures happen in these channels, and the probability of any eventual rupture. Figure 2(b) shows the survival curves (probability of no rupture in a channel) and the corresponding error intervals obtained by standard Kaplan-Meier survival analysis [18]. By intuition, we would expect that larger channels would have more ruptures and ruptures occur sooner because more cells enter the larger channels—meaning more possible cells to dissociate. Surprisingly, the survival curves are—to the level of our statistical uncertainties—independent of channel width. We also see that the survival probability saturates to ∼30% at long times, i.e., roughly ∼70% of channels have a dissociation event within the experimental measurement.

To see the correlation between cluster size and rupture time, we make a scatter plot of cluster size as a function of the rupture time for the 50-micron channels; each dot in Fig. 2(c) corresponds to one dissociation event [analogous plots for smaller channel sizes shown in Fig. S1(b)]. Roughly 50% of the dissociations are single-cell ruptures; the dots corresponding to these are primarily in the bottom-left corner of Fig. 2(c), showing these single-cell events often occur at early times. On the other hand, larger ruptures are relatively infrequent and tend to occur at later times. This trend is further supported by binning the data into small time ranges, where we can see that the average cluster size—black circles in Fig. 2(c)—increases with rupture time. We believe the average cluster size increases with time in part simply because to have a rupture of *n* cells, we must wait for *n* cells to enter into the channel.

## II. MODEL

### A. Phase field approach

To capture the complex shapes of cells, we use a multi-cell phase field approach [19–22] to track the cell boundaries. We can model a cell interface by introducing an auxiliary phase field *ϕ*_*i*_(**r**) for cell *i*. This field is *ϕ*_*i*_(**r**) = 1 inside the cell and *ϕ*_*i*_(**r**) = 0 outside cell *i*. Consequently, the cell interface can be easily tracked at *ϕ*_*i*_(**r**) = 1*/*2 [22]. From a physical perspective, this coarse-grained auxiliary field should be on a mesoscopic scale, much larger than the molecular structure inside the cell, but still smaller than the system size.

To model the dynamics of the cell, we adopt a Newtonian-like advection scheme in an overdamped environment [7, 23]:

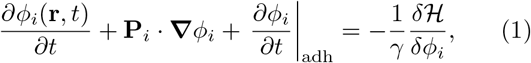

where the first two terms on the left-hand side describe cell *i* as moving with a velocity **P**_*i*_—i.e., the field is advected with velocity **P**_*i*_. The third term describes the evolution of *ϕ*_*i*_ due to cell-cell and cell-wall adhesion, which we will describe in detail later. On the right-hand side we introduce the functional derivative of a Hamiltonian ℋ [9, 23, 24]. This term tends to minimize ℋ [22]. *γ* is a friction coefficient so the whole Eq. (1) can be thought of as a force balance in an overdamped environment [25]—though see note [26]. We include a Cahn-Hilliard term, an area constraint, and cell-cell and cell-wall exclusion in the Hamiltonian [7, 23]:

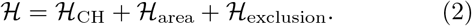

The first term in ℋ is the Cahn-Hilliard energy [9, 23]

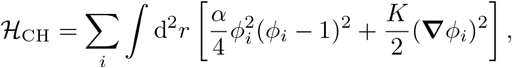

where 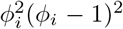 in the integrand is a double-well potential ensuring there exist two minima corresponding to the two phases *ϕ*(**r**) = 0 and *ϕ*(**r**) = 1; the second term (***∇****ϕ*_*i*_)^2^ tells the energy cost of forming a domain wall between the two phases. In general, the Cahn-Hilliard energy describes the interfacial tension [27] and controls the interfacial width 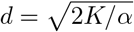 (see Appendix C). Also, we assume our cells to resist changes in area, introducing a term [9, 23]

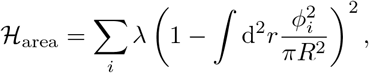

which penalizes any deviation from the preferred area *πR*^2^. We next introduce an energetic penalty to prevent cells from overlapping with other cells [23] or the walls of the microchannel,

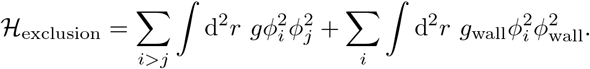

The first term in ℋ_exclusion_ describes cell-cell exclusion, which is a penalty for any overlapping between cells, since the integrand is only non-zero when both *ϕ*_*i*_ and *ϕ*_*j*_ are non-zero; the second term describes cell-wall exclusion in the same way as cell-cell exclusion—*ϕ*_wall_(**r**) is another auxiliary field introduced to depict the geometric confinement—*ϕ*_wall_ = 0 in the microchannel’s walls and *ϕ*_wall_ = 1 outside.

The term *∂ϕ*_*i*_*/∂t* |_adh_ in Eq. (1) models the effect of cell-cell and cell-wall adhesion,

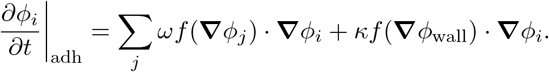

The first term treats cell-cell adhesion with the advection-like term of [21], and describes the attraction interactions between interfaces of different cells. Note that ***∇****ϕ* is a vector pointing normally into the cell described by *ϕ*. 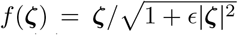is a function that saturates at large |***ζ***| to avoid numerical instability [21]. We have chosen this form over alternate implementations [20, 28], as we have found it possible to resolve high cell-cell adhesions without numerical instability. The second term in *∂ϕ*_*i*_*/∂t* |_adh_ models the cell-wall adhesion in the same way, with a strength *κ*.

Each cell in our model has a polarity **P**_*i*_ = *p*_*i*_(cos *θ*_*i*_, sin *θ*_*i*_); the polarity is the velocity the cell would have in the absence of any interactions with other cells or obstacles. We have set the default value of the magnitude *p*_*i*_ to *p*_0_ = 13.3 *μ*m*/*h—at the typical experimental cell velocity scale, though slower than cells that typically break off [16]. For the direction *θ*_*i*_, we assume that cells tend to align their polarity to the direction *θ*_0_ of the chemical gradient but there are still fluctuations that would lead to rotational diffusion,

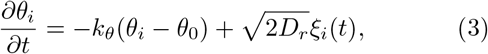

where *ξ*_*i*_(*t*) is a Gaussian white noise satisfying ⟨*ξ*_*i*_(*t*) ⟩ = 0, and ⟨*ξ*_*i*_(*t*)*ξ*_*j*_(*t*^*′*^)⟩ = *δ*_*ij*_*δ*(*t* −*t*^*′*^), and the preferred direction *θ*_0_ is determined by the chemotactic cue. The first term on the right-hand side drives *θ*_*i*_ to relax to *θ*_0_ on a timescale of ∼ 1*/k*_*θ*_. In the absence of chemotaxis (*k*_*θ*_ = 0), the angle *θ*_*i*_ will simply diffuse with angular diffusion coefficient *D*_*r*_. This representation of cell polarity is a very simplified one, and could be extended by models of Rho GTPase dynamics [17, 29] or stochastic protrusion dynamics [30].

### B. Cell states—chemotaxis and leader cells

Above, we described cells as following a chemoattractant gradient. However, not all of the cells are likely to be able to sense the chemical signal—since the chemoattractant concentration will be diluted as it reaches the bottom of the microchannel. Rather than explicitly modeling chemoattractant dynamics, we make a simple assumption and distinguish the cells that are in the seeding region (reservoir at the bottom of Fig. 1), and cells that are in the microchannel. We assume cells located in the seeding region are not guided by the chemical gradient—cells in the seeding region are just followers of those guided cells moving along the chemical gradient in the channel.

In addition, there is well-established evidence that in collective migrations like wound healing [1], and cancer metastasis [4, 5], cells invade the free space under the apparent guidance of some high motile “leader” at their leading edge [12, 31], which can be crucial in collective invasion [32, 33]. These leader cells can invade the free surface more easily than other cells and coordinate their motion with their followers. Within cancerous collectives, leader cells may be keratin-14 positive cells which can either activate at or move to the leading edge and guide collective migration in tumor invasion [34–36]. We include leader cells within our model—these are cells at the front that have a larger self-propulsion strength *p*_*i*_ than other cells.

We illustrate the three cell states—follower (green), guided (yellow), and leader (purple)—described in our model in Fig. 3. The blue arrows represent the polarity **P**_*i*_ and the red arrows indicate the center of mass velocity 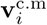. (see Appendix D). Followers (green) are cells with chemotaxis strength *k*_*θ*_ = 0 and polarity magnitude *p*_follower_ = *p*_0_. Guided cells (yellow) sense the chemoattractants and typically exhibit higher velocities than followers—modeling chemokinesis and chemotaxis [37]— i.e., guided cells have a non-zero *k*_*θ*_ and larger magnitude of polarity *p*_guided_ = 2*p*_0_. At the time a cell reaches the front, it has a probability of becoming a leader cell is dependent on its contact length with other cells:

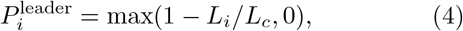

where *L*_*i*_ is the contact length of the *i*th cell with its neighbors, and *L*_*c*_ is a characteristic length. In practice, we count the number of cell-cell contact points *n*_*i*_ for each cell, hence 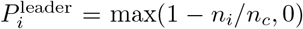, where *n*_*c*_ is the characteristic number of contact points (see Appendix D for details). Eq. (4) reflects the idea that cells that cell-cell contact inhibits leadership—similar to the proposal of [12].

**FIG. 3.**
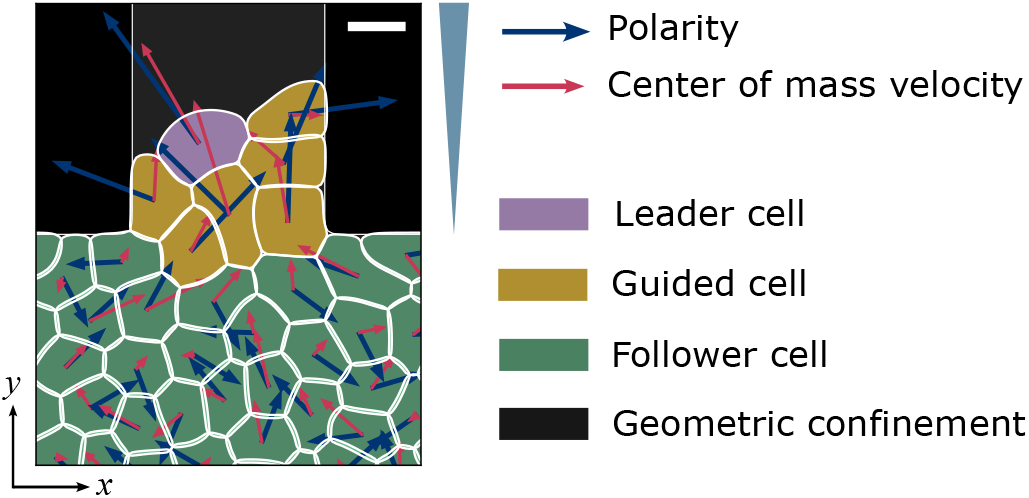
Schematic of the model. For each cell, the blue arrow represents its polarity **P**_*i*_, and the red arrow shows its center of mass velocity 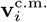 ; chemoattractant concentration increases in the +*y* direction. Scale bar, 15 *μ*m.

The only distinction between leader cells and guided cells is that leader cells have stronger self-propulsion *p*_leader_ = 3*p*_0_ (see Appendix D for details of how we define reaching the front in the model). This is consistent with our earlier experimental measurement [16], where cells that eventually dissociate in 6-micron channels can have larger speeds (around ∼ 30 *μ*m*/*h).

Currently, the best model we find that can fit the experimental data (as shown in Fig. 2) is composed of these three types of cells: Cells near the channel entrance and inside the channel are identified as guided cells (labeled yellow in Fig. 3), and cells in the chamber are the followers (green) following the guided cells, while leader cells (purple) are tip cells at the invading front. Alternate models without leader cells are shown in Appendix A.

## IV. MODEL RESULTS

### A. Matching with experimental results

We simulate our phase field model for 18 hours, after an initial equilibration time where cells relax to more physical shapes (see Appendix D). Figure 4 shows the simulation snapshots for *W* = 6, 10, 20, 50 *μ*m microchannels (see also Supplemental Movies 1–4). We observe both single-cell and cluster ruptures. Figure 5 shows statistics on rupture size and times for our simulations, analogous to our experimental results in Fig. 2. We have been able to recapitulate several key factors of the experiment. Specifically, our model reproduces (i) the predominance of single-cell ruptures, (ii) the occurrence of larger ruptures in wider channels, (iii) the independence of survival curves from microchannel width, and (iv) the increase of average cluster size with rupture time.

When varying the channel width, we include the possibility that cell tension and adhesion dynamics change in confinement, as discovered earlier [16, 38, 39]. This unavoidably requires some degree of tuning of parameters as a function of channel width. In particular, to match the survival probability with experiments as shown in Fig. 2(b), we have to increase cell-cell adhesion strength *ω* from 0.7*ω*_0_ to *ω*_0_ for channel widths from 6–50 *μ*m, where *ω*_0_ is the reference value of adhesion strength in our model (see Table I for details). This is a relatively small tuning of adhesion strength, but is sufficient to ensure that the survival curves do not differ strongly as a function of channel width [Fig. 5(b)]. If we do not make this assumption, and instead assume that all channels have the same value of cell-cell adhesion strength *ω*_0_, we find that the 10-micron channel has slower rupture than other channels [Figs. 10(b) and 10(f)]. This is because cells in the 10-micron channel have a broader cell-cell contact line than the 6-micron channel—but in the 20-micron channel, multiple cells can enter at once, so ruptures do not require breaking a larger contact line (see Appendix B and Fig. 11).

**TABLE I.** Table of simulation parameters^a^.

| Parameter | Description | Dimension | Dimensionless value | Real value |
| --- | --- | --- | --- | --- |
| $\delta x$ | Lattice constant | $L$ | 1 | $1\ \mu\text{m}$ |
| $\delta t$ | Simulation time unit | $T$ | 1 | 15 s |
| $N$ | Number of cells | 1 | 48 | 48 |
| $\Delta t$ | Simulation time step | $T$ | 0.4 | 6 s |
| $R$ | Target cell radius | $L$ | 7 | $7\ \mu\text{m}$ |
| $p_0$ | Advection velocity (polarity) | $L/T$ | $5.556 \times 10^{-2}$ | $13.3\ \mu\text{m/h}$ |
| $D_r^{-1}$ | Reciprocal of the rotational diffusion coefficient | $T$ | 180 | 45 min |
| $d$ | Interfacial thickness | $L$ | 2 | $2\ \mu\text{m}$ |
| $\gamma$ | Friction coefficient | $ET/L^2$ | 12 | |
| $K$ | Cell tension coefficient | $E$ | 3.5 | |
| $\lambda$ | Cell area coefficient | $E$ | 250 | |
| $g$ | Cell-cell repulsion coefficient | $E/L^2$ | 4 | |
| $g_{\text{wall}}$ | Cell-wall repulsion coefficient | $E/L^2$ | 5 | |
| $\omega$ | Cell-cell adhesion coefficient | $L^2/T$ | 3.5–5 | |
| $\omega_0$ | Reference value of adhesion coefficient | $L^2/T$ | 5 | $\sim 0.3\ \mu\text{m}^2/\text{s}$ |
| $\kappa$ | Cell-wall adhesion coefficient | $L^2/T$ | 3.5 | $0.23\ \mu\text{m}^2/\text{s}$ |
| $\epsilon$ | Regularization coefficient | $L^2$ | 37.25 | $37.25\ \mu\text{m}^2$ |
| $k_\theta$ | Chemotaxis strength coefficient | $1/T$ | $10^{-3}$–0.05 | $0.004$–$0.2\ \text{min}^{-1}$ |
| $k_\theta^0$ | Reference value of chemotaxis strength coefficient | $1/T$ | 0.0025 | $0.01\ \text{min}^{-1}$ |
| $n_c^0$ | Default characteristic number of contact points $n_c$ | 1 | 256 | |

<sup>a</sup> These parameters are used throughout the paper; any deviation from them is explicitly noted.

**FIG. 4.**
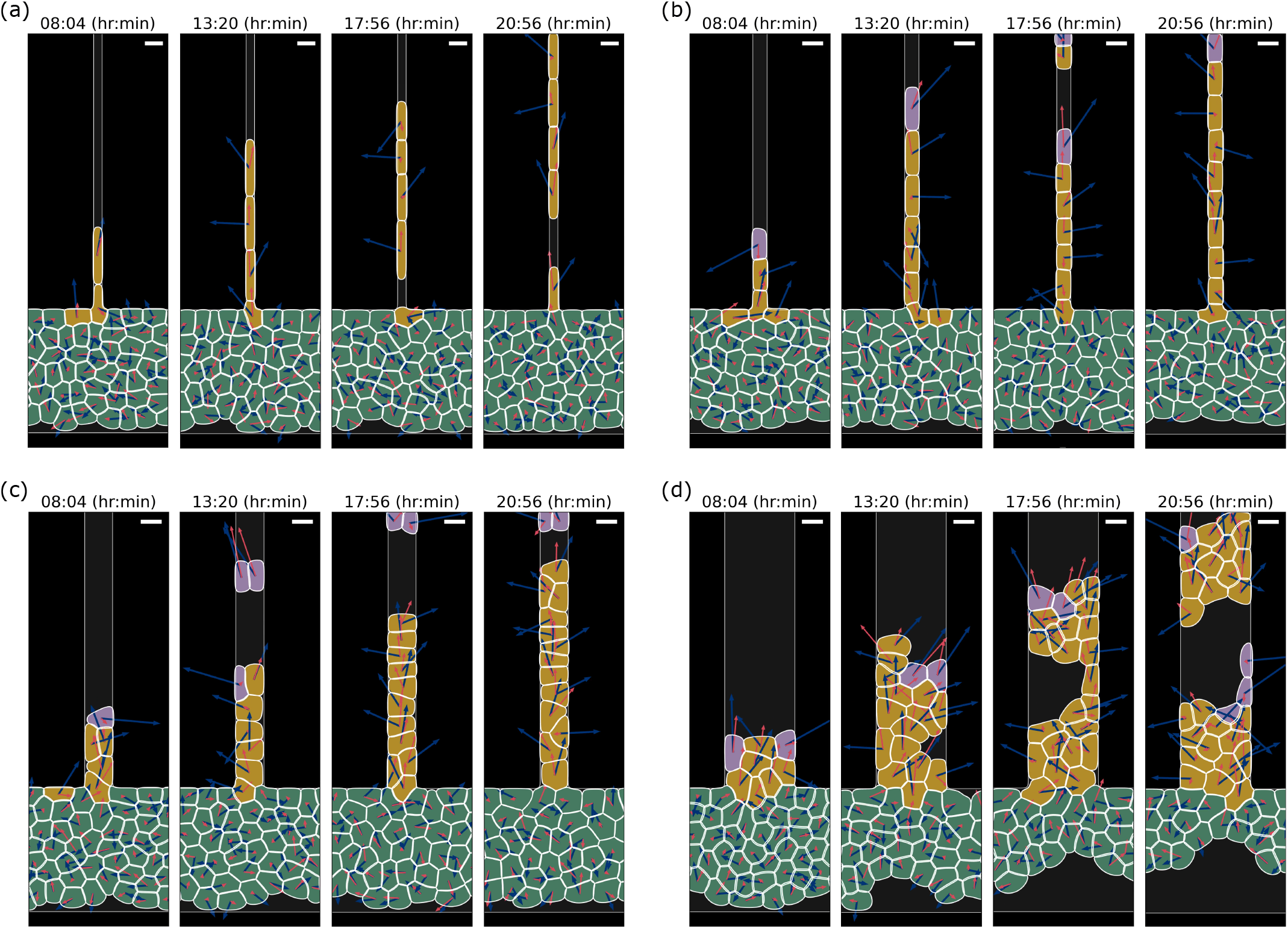
Evolution of rupture for simulations of cells in microchannels with different widths. Panels (a)–(d) correspond to *W* = 6, 10, 20, and 50 *μ*m, respectively. There are chemical gradients along the microchannels (+*y* direction) driving cells to move upwards. Scale bars, 15 *μ*m.

**FIG. 5.**
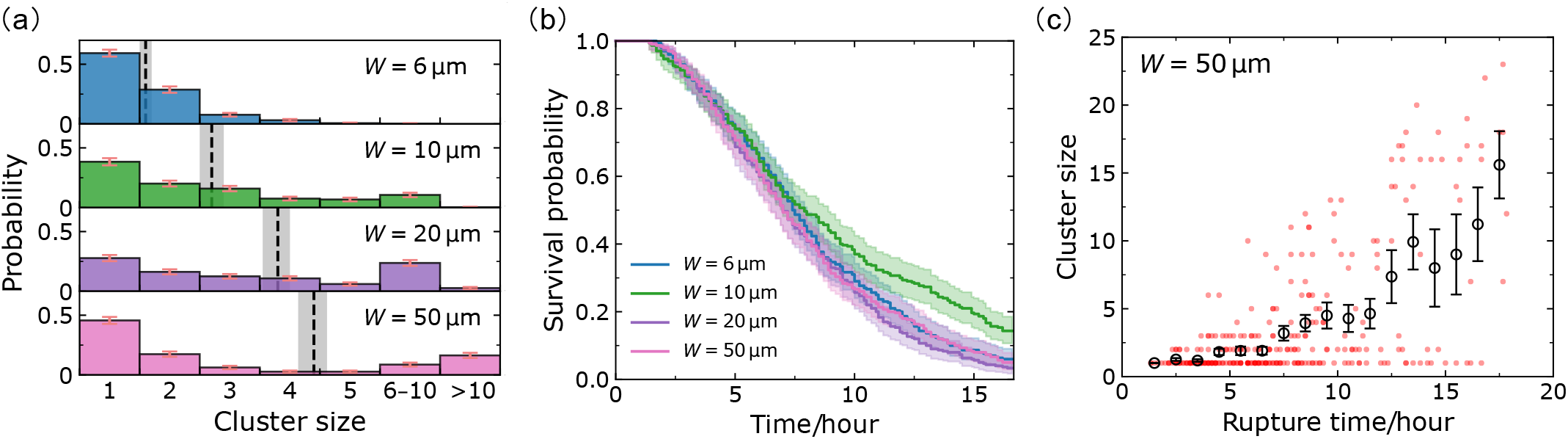
Simulation results to match experimental data. We adjust *ω* = 0.7*ω*_0_ for the two narrowest channels, *ω* = 0.8*ω*_0_ for 20 *μ*m channels, and *ω* = *ω*_0_ for 50 *μ*m channels. (a) Cluster-size distributions for *W* = 6, 10, 20, 50 *μ*m microchannels. The corresponding average cluster sizes (dashed lines; mean *±* SE) are *A* = 1.6, 2.7, 3.8, 4.4. The complete histograms are shown in Fig. S2(a). Panel (b) shows the survival probabilities for different channel widths; the final survival probabilities are *K*_*s*_ = 16*/*300, 43*/*300, 8*/*300, 13*/*300, respectively. Panel (c) shows the (rupture time, cluster size) phase plane scatter plot. The circles represent the average cluster sizes within each time bin (1 hour), and the error bars indicate the corresponding standard errors. Figures for all channel widths are given in Fig. S2(b). We conduct 300 independent simulations for each microchannel width *W* with 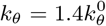 in all channels; *ω*_0_ and 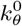 represent the reference values for adhesion and chemotaxis strength in the simulation.

The experimental cluster-size distributions [Fig. 2(a)] are well captured by our model [Fig. 5(a)]. We observe that single-cell ruptures are always predominant, though there is a population of larger ruptures in wide channels, leading to a mean rupture size that increases with channel width. We note that the single-cell ruptures being predominant depends on the presence of leader cells; we show how cluster statistics depend on the leader-cell number in Sec. IV F, and show results without leader cells in Figs. 10(c) and 10(d).

We plot rupture cluster size as a function of rupture time in Figs. 5(c) and S2(b). Similar to Fig. 2(c), we find that small ruptures dominate and usually happen at early times, and large ruptures are less probable and tend to happen at later times. Moreover, we see—consistent with the experiment—that the average cluster size increases with rupture time.

We note that the survival curves in Fig. 5(b) drop nearly to zero, which the experimental curves do not. We think this may reflect the evolution of parameters like the chemoattractant strength and cell-cell adhesion over time in our experiments, and address this further in Sec. V.

Next, we systematically adjust several key parameters in the model to investigate their impact on the rupture behavior of the monolayers, including the cell-cell adhesion strength *ω*, cell-wall adhesion strength *κ*, chemotaxis strength *k*_*θ*_, and the probability for tip cells to become leader cells.

### B. Rough intuitive picture: active escape over a barrier

Thinking about a chain of invading cells, such as shown in Fig. 4, we expect that rupture is not likely to happen if all cells are moving with consistent velocities. Differences in the self-propulsion from one cell to another lead to the cell centers moving apart from one another—increased strain. This difference in self-propulsion could arise because cell polarities point in different directions, e.g., due to the rotational noise *D*_*r*_, or have different magnitudes, due to one cell having a different state from its neighbors. The difference in motilities between cells tends to work against forces keeping cells together—largely cell-cell adhesion. In this way, we think of rupture as being initiated by fluctuations of cell motility leading to the cells crossing an energy barrier [40]. For channels wider than a single cell can span, a dissociation event will start with the breakage of a single cell-cell junction, but this fracture will then have to evolve to cross the channel, as seen in, e.g., Refs. [41, 42] and in Fig. 1(b) for the 50-micron channel. Our picture here is an oversimplification because there is not a single energy barrier, but a complex process involving re-arrangement of cells and cell boundary deformations. Nonetheless, this rough picture makes some clear predictions. We would expect faster rupture if the cell motion is noisier, and more rupture if there is more variability in cell speeds—both increasing the tendency for cells to be pulled apart. We would expect less rupture if there is high cell-cell adhesion (increasing the barrier height)—and if the process of rupture is nucleated at the walls, then cell-wall adhesion could also influence the barrier. We test all of these ideas below.

### C. Cell-wall adhesion allows cells to invade into narrow channels

Cell-wall adhesion is vital to observe ruptures in narrow channels. Figures 6(a) and 6(b) show the clustersize distributions and survival curves when we turn this term off. The rupture behavior in wide channels (like 50 *μ*m channels) is not immediately different in the absence of cell-wall adhesion, but ruptures are strongly suppressed in narrow (6 *μ*m or 10 *μ*m) channels—most dramatically, no ruptures occur in 6 *μ*m microchannels. This is because, in this scenario, cells fail to enter the narrow channels (see Supplemental Movie 5). We change the cell-wall adhesion strength *κ* and see how this influences rupture in narrow (6-micron) and wide (50-micron) channels. As shown in Fig. 6(c), *κ* has little effect on 50 *μ*m channels, but it can control the rupture probability (1 −final survival probability) in narrow channels. Rupture fraction in small channels increases rapidly with cell-wall adhesion and then saturates, consistent with the idea that cell-wall adhesion is helping cells overcome an energy barrier at the channel entrance [43]. The rupture fraction is low at small cell-wall adhesion because in these cases, few cells manage to enter the channel. We plot the probability of any cells entering the channel in Fig. 6(c) using dashed lines, and it closely tracks the rupture fraction—at small *κ*, if any cell enters the channel, there will almost always be a dissociation. Interestingly, rupture probability in both 6- and 50-micron channels eventually decreases as cell-wall adhesion is increased. We believe this is because some ruptures require detachment from the wall as part of their dissociation process—e.g., the formation of the “notch” in the 50-micron channel simulation in Fig. 4(d) and Supplemental movie 4. Cell-wall adhesion thus has two effects: some amount of cell-wall adhesion helps cells enter channels, but large cell-wall adhesion helps prevent cells from dissociating.

**FIG. 6.**
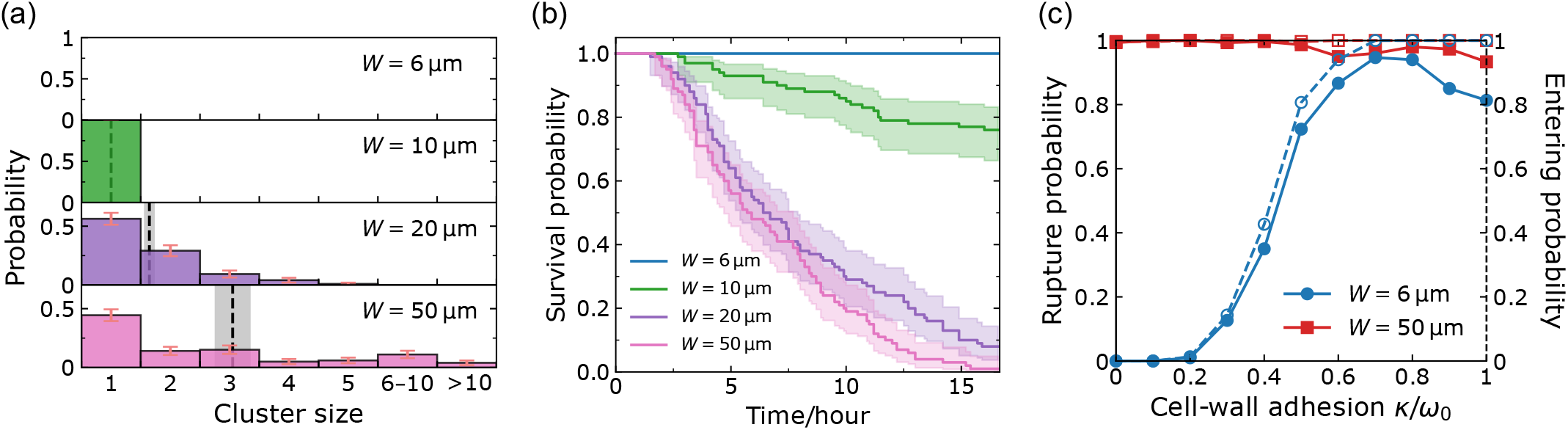
Cell-wall adhesion helps cells invade into narrow channels. Panels (a) and (b) are simulation results (100 independent simulations) when there is no cell-wall adhesion term. The corresponding average cluster sizes (dashed lines; mean *±* SE) are *A* = 0, 1.0, 1.6, 3.1, and the final survival probabilities are *K*_*s*_ = 100*/*100, 76*/*100, 8*/*100, 1*/*100. The complete histograms are shown in Fig. S3(a). Panel (c) shows the rupture probability 1 −*K*_*s*_ (solid lines) and the probability of cells entering the channel (dashed lines) as a function of cell-wall adhesion strength *κ* for narrow (6 *μ*m) and wide (50 *μ*m) microchannels; 300 independent simulations for each point.

### D. Strong cell-cell adhesion leads to rarer but larger ruptures

Ruptures occur at cell-cell junctions for epithelial monolayers, and the intercellular adhesion complexes, such as different members of the cadherin family, can control the strength of tissues [41]. In our model, cell-cell adhesion strength *ω* is one of the most important parameters that prescribe the rupture behavior. Our intuition is that increasing cell-cell adhesion strength *ω* should increase the barrier to rupture and reduce the frequency of rupture events. Figures 7(a) and 7(b) show the simulation results for 50 *μ*m microchannels. With the increase of cell-cell adhesion, the survival curve drops much more slowly, and the final survival rate rises a little—rupture is less common due to the strong connection between cells. This influence of cell-cell adhesion is consistent with our earlier experimental work, which found that in 6-micron channels, increasing the cell-cell adhesion by overexpressing E-cadherin led to suppressed rupture, and downregulation by shRNA led to increased rupture [16].

**FIG. 7.**
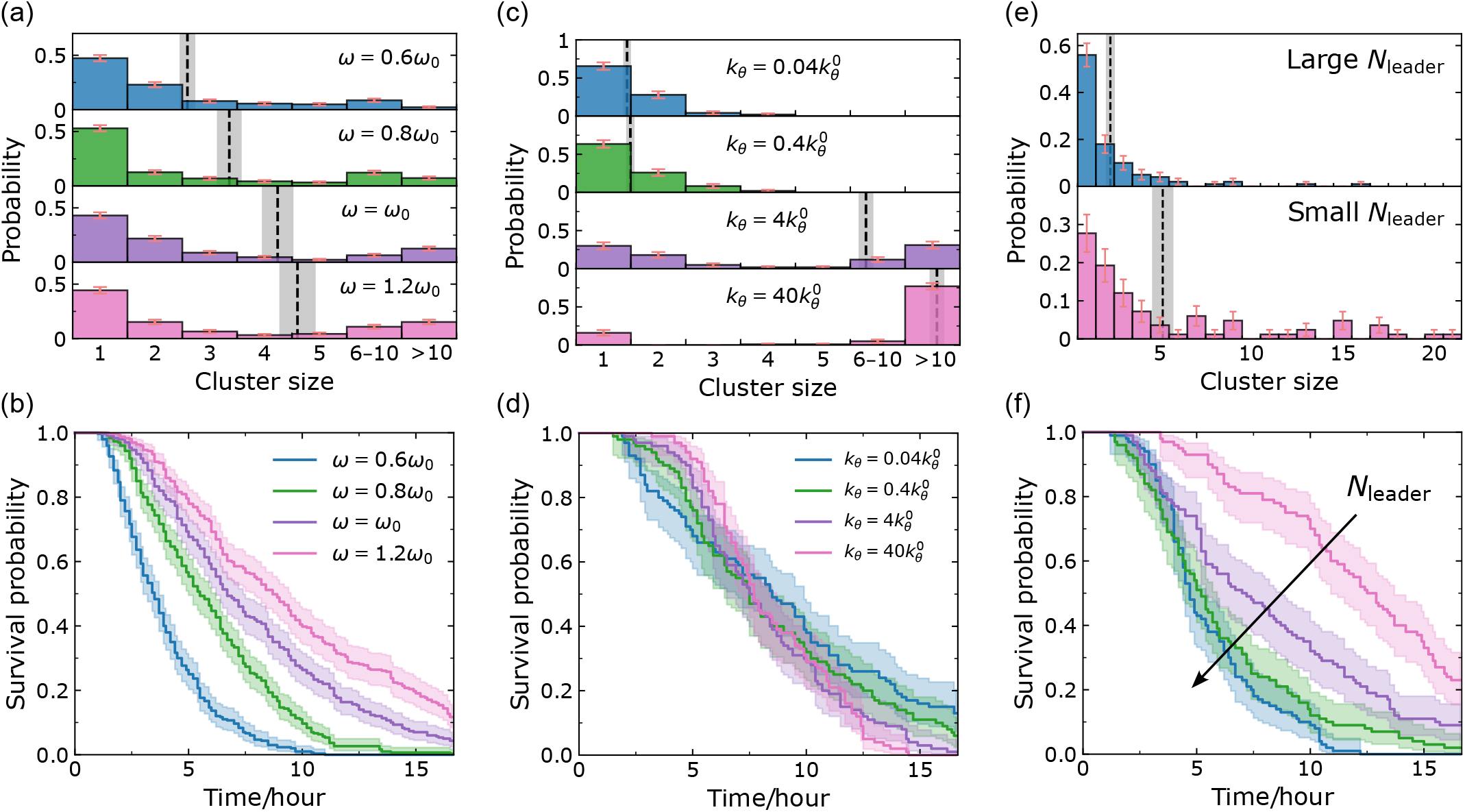
How intercellular adhesion, chemotaxis, and leader cells affect the dissociation behaviors. Panels (a), (b) are simulation results for different cell-cell adhesion strength *ω* in *W* = 50 *μ*m channels. The corresponding average cluster sizes (dashed lines; mean *±* SE) are *A* = 2.6, 3.4, 4.2, 4.6, and the final survival probabilities are *K*_*s*_ = 0, 1*/*100, 4*/*100, 11*/*100. Panels (c), (d) are simulation results for different chemotaxis strength *k*_*θ*_ in *W* = 50 *μ*m microchannels. The corresponding average cluster sizes (dashed lines; mean *±* SE) are *A* = 1.4, 1.5, 6.9, 13.4, and the final survival probabilities are *K*_*s*_ = 12*/*100, 6*/*100, 1*/*100, 0*/*100. Panels (e), (f) are simulation results when the probability of leader cell formation [Eq. (4)] is altered by setting the characteristic number of contact points *n*_*c*_ to 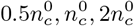, and 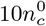, where 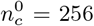 (see Appendix D). Larger *n*_*c*_ corresponds to larger *N*_leader_. The corresponding average cluster sizes (dashed lines; mean ± SE) are *A* = 2.3, 5.2. The complete histograms for panels (a), (c) are shown in Fig. S3(b), (c).

From the cluster-size distribution, we find there are more ruptures of larger clusters (*>*10 cells) happening in systems with strong cell-cell adhesion, and mean cluster sizes increase with increasing adhesion *ω*. In general, strong cell-cell adhesion leads to less rupture, but if we get a rupture, it is more likely to be a large rupture. Our results show that the influence of changes in cell-cell adhesion on metastasis may be complex, and we would expect its effect on overall risk to depend on the relative importance of the rate and size of metastases, even neglecting any effect of cadherins on cell proliferation and survival.

### E. Strong chemotaxis strength leads to larger and faster ruptures

We expect that the rate of rupture could potentially be increased if the cells are noisier in their motility—i.e., if they are less uniformly guided by the presence of the chemoattractant gradient. This is in part controlled by the chemotaxis strength *k*_*θ*_.

To better understand the magnitude of *k*_*θ*_, we think about the dynamics of cell orientation *θ* for a single isolated cell, which follows Eq. (3). We take the preferred direction *θ*_0_ = 0 then by the properties of Ornstein-Uhlenbeck processes [44], the average is ⟨*θ*⟩ = 0, and the variance is ⟨*θ*^2^⟩ = *D*_*r*_*/k*_*θ*_. Cell chemotactic accuracy is often characterized by a “chemotactic index” CI = ⟨cos *θ*⟩ (or related definitions) [45]; CI = −1 would be perfect chemorepulsion, CI = 0 is undirected migration, and CI = 1 is perfect, deterministic motion up the gradient. For our model, we can compute for an isolated cell CI = ⟨cos *θ*⟩ = exp(−⟨*θ*^2^⟩ */*2) = exp(−*D*_*r*_*/*2*k*_*θ*_), using the Gaussian distribution of *θ*. This gives us a way to interpret *k*_*θ*_. Our default value of *k*_*θ*_ = 1.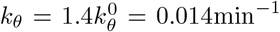 corresponds to a chemotactic index of ⟨cos *θ*⟩ ≈0.45. However, this estimate—which essentially assumes an isolated cell not in confinement—may not always reflect the chemotactic accuracy of cells in our full simulation. First, in our system, we have many cells—so the motion of their center of mass will be more accurate than any individual cell [45]. Secondly, cells in narrow channels are forced to have their velocity in the direction of the channel, so they will be more directional in confinement [13, 46].

Figures 7(c) and 7(d) show the cluster-size distributions and the survival curves for different chemotaxis strengths *k*_*θ*_. These simulations are in 50 *μ*m microchannels, and cell-cell adhesion strength is set to the default *ω* = *ω*_0_. We vary *k*_*θ*_ across a large range (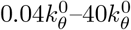, which corresponds to a CI range of roughly 0–1). From Fig. 7(c), we see that more ruptures of larger size happen at large *k*_*θ*_ when cell chemotaxis is more deterministic— cells move together as a large pack. In Fig. 7(d), when there is only very weak guidance (small *k*_*θ*_), when the cells can be polarized in almost any direction, not only toward the channel, we can see more early ruptures (survival curve drops earlier) but also see more channels that never have ruptures. Removing chemoattractant entirely in the experiments of [16] did reduce ruptures significantly. By contrast, at very strong guidance (large *k*_*θ*_), the survival curves become steeper, indicating that a large chunk of cells move together and faster, and large clusters are the majority outcome.

### F. Leader cells drive single-cell rupture

Within our model, the key factor that controls whether cells have variable states is the probability for cells that reach the leading edge to become “metabolically active” (high-speed) leaders, given by Eq. (4). We can increase the probability of cells becoming leader cells—and thus the average number of leader cells *N*_leader_—by increasing the characteristic number of contact sites *n*_*c*_, and vice versa.

Figure 7(e) shows the cluster-size distributions for large *N*_leader_ and small *N*_leader_, and Fig. 7(f) shows the survival curves for different *N*_leader_ in 50 *μ*m channels, with *ω* = *ω*_0_, and 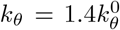. The survival curves decrease faster with larger *N*_leader_, which tells us that more leader cells lead to faster ruptures. Also, from Fig. 7(e) we can see that the leader cells increase the ratio of single-cell ruptures to all ruptures, which ensures the absolute predominance of single-cell ruptures shown in experiments. These leader cells, which have the largest velocity in the system and are located at the leading edge of the expanding monolayer, can dissociate from the main bulk more easily. Since leaders are relatively rare in our default parameters, this may explain the universal most probable outcome of single-cell rupture in all cluster-size distributions.

### G. Unjamming is necessary but not sufficient to create ruptures

A confluent monolayer can go from a jammed state to an unjammed state, allowing cells within the monolayer to more freely rearrange [47, 48]. Experiments have shown that jamming is influenced by factors such as cell shape, cell motility, and cell-cell adhesion [48–50], and the spatiotemporal control of tissue states may represent a generic physical mechanism of embryonic morphogenesis [51]. Ref. [7] shows that several different order parameters can capture this transition behavior. For instance, if a dimensionless quantity, called shape index, 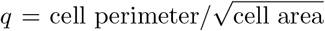, is larger than some critical value *q*_*c*_ (Ref. [8] indicates *q*_*c*_ ≈ 3.81), then the system is in a fluid-like unjammed state, where cells can squeeze between neighbors; if *q* is smaller than *q*_*c*_, the system is in a more solid-like jammed state, where cells are somewhat caged by their nearest neighbors. Refs. [7] and [8] also show that a system in the solid state can be fluidized by increasing cell motility, cell deformability, or the preferred shape index of the tissue.

Does the jamming transition influence the presence of rupture? To characterize the extent of jamming, we follow [7] and measure the effective diffusivity of cells as the order parameter, 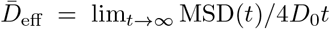, where MSD(*t*) is the mean squared displacement of the cells (see Appendix D). Here, 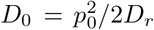 is the diffusion coefficient for an isolated follower cell [7, 8]. We control jamming by varying cell-cell adhesion and cell speed *p*_0_. To compare with previous work, we characterize the cell’s speed by computing the dimensionless Péclet number Pe = *p*_0_*/*(*RD*_*r*_) [7], which is roughly speaking the distance an isolated cell would travel in the time the cell takes to reorient (1*/D*_*r*_), scaled by its size *R*. We measure the effective diffusivity of cells in the rectangular confinement (with periodic boundary condition in the *x* direction, see Fig. 8) of our previous simulations—but don’t open the channel entrance, so all cells are confined in the original reservoir and there is no invasion. As shown in Fig. 9(a), we see a transition between near-ballistic motion at short time scales to more-diffusive motion at long times (*t >* 1*/D*_*r*_). Mean squared displacements are naturally larger for cells with larger motility *p*_0_ (larger Pe). Fitting to a linear function to extrapolate to measure 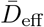, we can see that there is a clear transition in 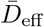 representing a fluid-solid transition [Fig. 9(b)]. Cells with low motility (the first two data points below the red dashed line) are in the solid-like jammed state, and cells with high motility (the rest points above the red dashed line) are in the unjammed state, which is consistent with Ref. [7, 8]. The red dashed line is the threshold value 0.012, which we use to indicate the presence of a solid-fluid transition. This value is chosen both based on the sudden transition in Fig. 9(b), but also on the appearance of simulation movies showing the degree of cell rearrangement. Simulation snapshots for a typical jammed state and unjammed state are given in Fig. 8. We see that cells in the jammed state are highly ordered and near-crystalline, while the unjammed state shows more elongated, disordered cell shapes, as in [7].

**FIG. 8.**
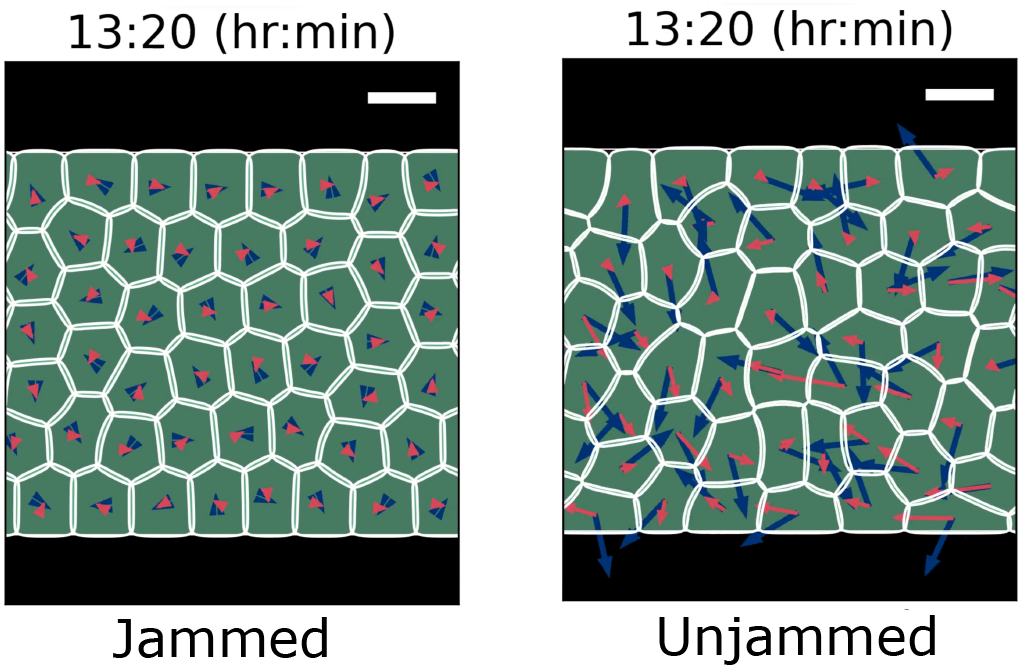
Simulation snapshots for typical systems in jammed (Pe = 0.14) and unjammed (Pe = 1.43) states at the same simulation time. Blue arrows represent the polarity **P**_*i*_ and red arrows indicate the center of mass velocity 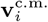.. Scale bars, 15 *μ*m.

**FIG. 9.**
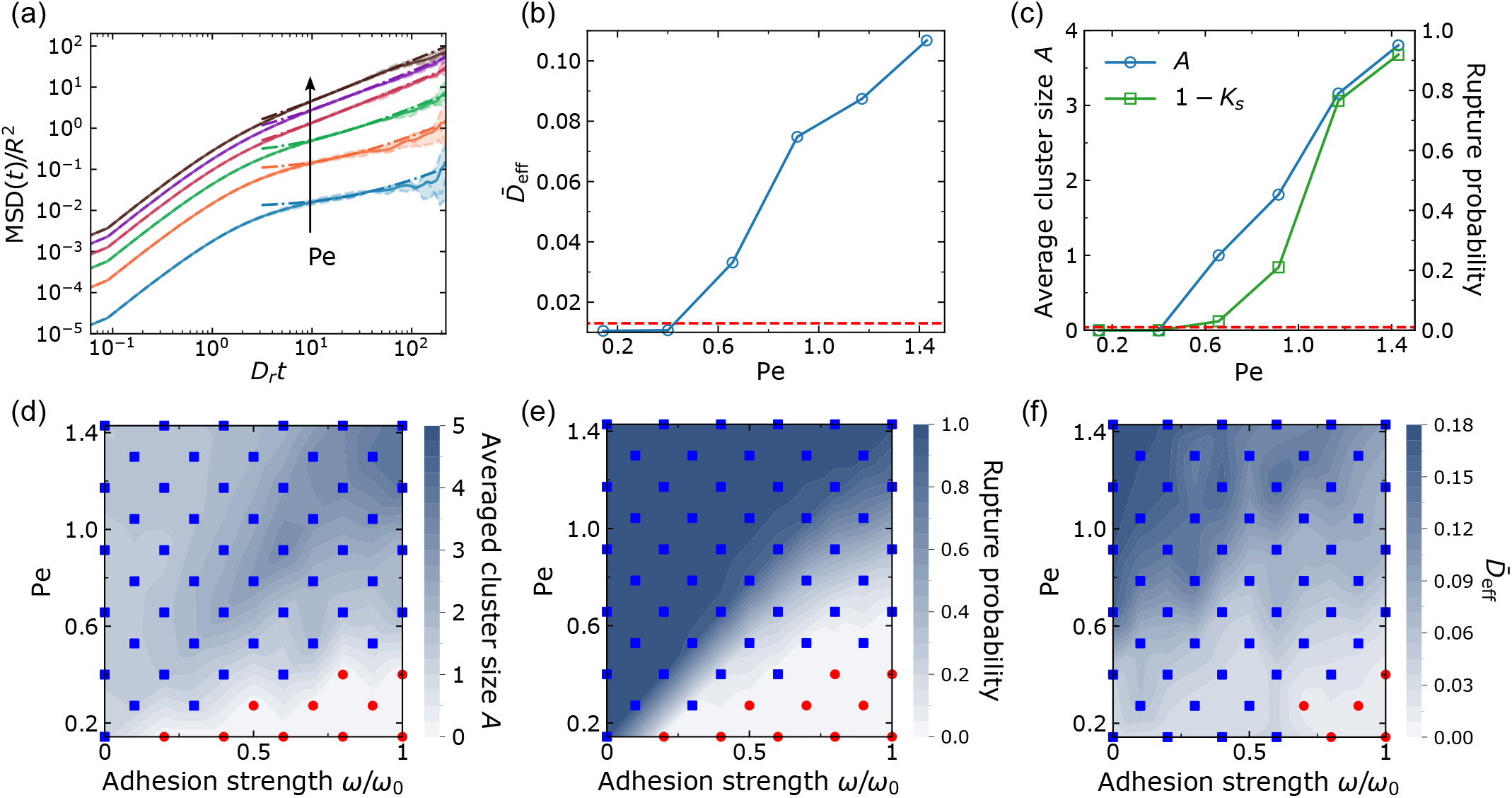
Unjamming and rupture. (a) Mean squared displacement MSD(*t*) for different cell motility quantified by a dimensionless Péclet number Pe := *p*_0_*/*(*RD*_*r*_). The different Pe shown in this panel are 0.14, 0.40, 0.66, 0.91, 1.17, and 1.43 (default); we have rescaled MSD and *t* to make them dimensionless; for each Péclet number, the dashed lines are results from three independent simulations, and the solid line is their average, and the dash-dotted lines are linear fit at late times where the effective diffusivity 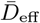 is extracted. (b) The effective diffusivity 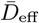 as an order parameter for solid-liquid transition. (c) Corresponding rupture transition of average cluster size (blue) and rupture probability (green). Panels (d) and (e) are phase diagrams for rupture transition. Red points indicate values equal to 0 and blue squares represent values greater than 0. (f) Phase diagram for jamming transition. Red points indicate values below the threshold value 0.012 and blue squares represent values above the threshold value. 300 independent simulations for each point in panels (c)–(f).

**FIG. 10.**
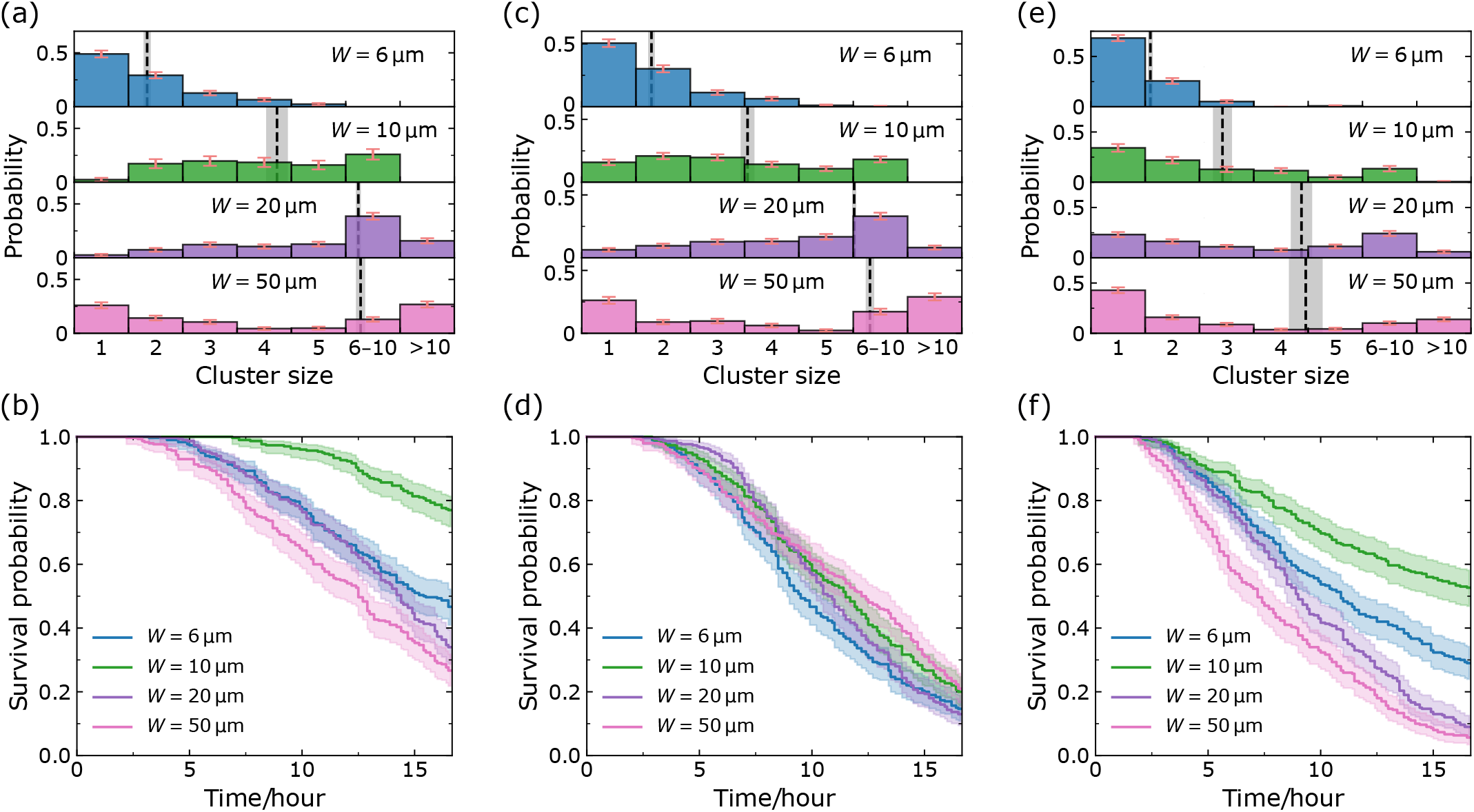
Simulation results for alternate models. Panels (a), (b) are simulation results with neither leader cells nor tuning of cell-cell adhesion (*ω* = *ω*_0_ in all microchannels). Panels (c), (d) are simulation results after we tune down cell-cell adhesion in narrow microchannels (but no leader cells). Panels (e), (f) are simulation results after we introduce the leader cells but still keep cell-cell adhesion *ω* = *ω*_0_ in all microchannels. 300 independent simulations; chemotaxis strength 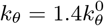.

**FIG. 11.**
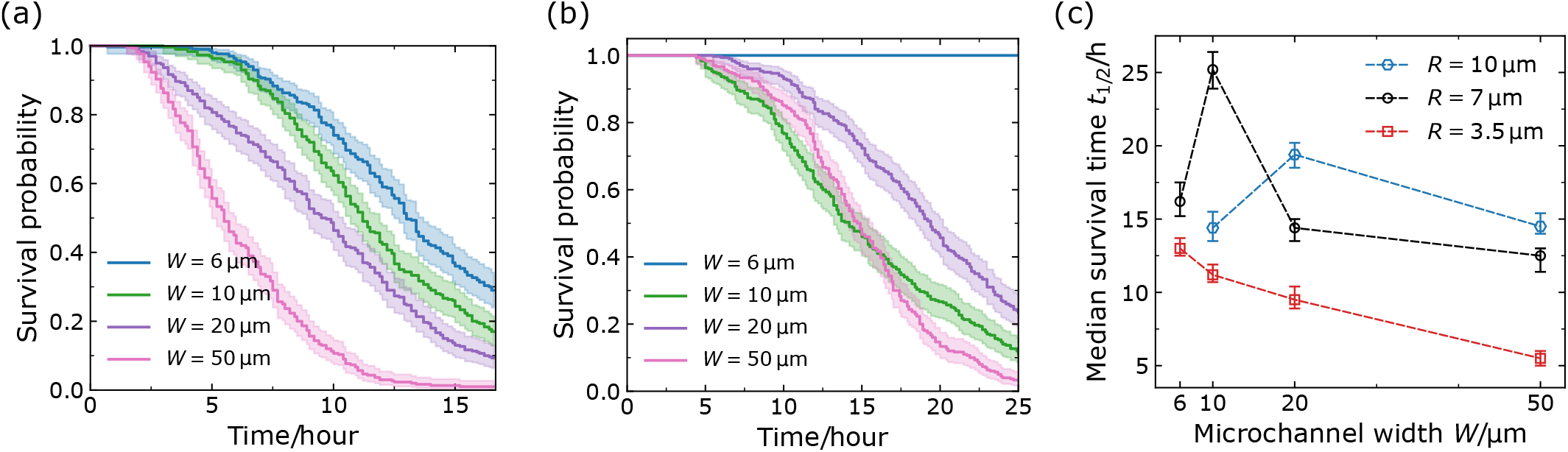
Simulation results for different cell sizes without special tunings (300 independent simulations, *ω* = *ω*_0_ in all microchannels, 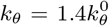, no leader cells). Panels (a) and (b) show the survival curves for different channel widths when the cell radius *R* = 3.5 *μ*m and 10 *μ*m, respectively. We notice that the 6-micron channels are too narrow for cells with a radius of 10 *μ*m to enter, resulting in no ruptures occurring [blue curve in Panel (b)]. Panel (c) shows the median survival time *t*_1*/*2_ as a function of microchannel width *W* for different cell sizes; the results for *R* = 7 *μ*m (black) are extracted from Fig. 10(b); the error bars indicate the 95% confidence intervals. When changing the cell radius *R* to 3.5 and 10 *μ*m, we proportionally adjust the interfacial thickness 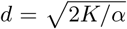 by varying *α* to fix *d/R*, and we also modify the number of cells *N* to 96 and 30 accordingly.

To study the relationship between this jamming transition and our dissociation statistics, we run simulations of ruptures in 50-micron channels using the same procedure we used for Fig. 5, but varying the Péclet number. While we are varying the motility Pe, we keep *R* and *D*_*r*_ constant and change the basal polarity magnitude to *p*_0_ = Pe · *RD*_*r*_. However, we still maintain the relationships *p*_guided_ = 2*p*_0_ and *p*_leader_ = 3*p*_0_, i.e., we are changing the self-propulsion of all cells in the system. Figure 9(c) shows there is no rupture for a system with *p*_0_ at its two lowest values—when our jamming simulations indicate the tissue is in a solid state—and ruptures start to happen after the system is fluidized by increasing cell motility.

Is this a coincidence, or is the jamming transition always aligned with the rupture transition? We sweep the (*ω*, Pe) parameter plane to construct several phase diagrams, as shown in Figs. 9(d) 9(f). Both Figs. 9(d) and 9(e) show the rupture transition and that ruptures are completely abolished when the tissue has strong cell-cell adhesion but lower motility (red-point regions in the bottom right). We also notice—consistently with our earlier results—that larger ruptures tend to happen in a system of cells with strong cell-cell adhesion and high motility [Fig. 9(d), upper right corner].

We also measure jamming as a function of adhesion and motility in Fig. 9(f). Consistent with prior research findings [7, 8], we have observed that a jammed state can be transitioned to an unjammed state by elevating cell motility. Furthermore, we also find the reinforcement of cell-cell adhesion will suppress unjamming, which is supported by experiments [14, 52].

Upon comparing Fig. 9(f) to Figs. 9(d) and 9(e), it is evident that the red-point region indicating the solid states in Fig. 9(f) is a subset of the red-point region corresponding to no rupture in Figs. 9(d) and 9(e). In other words, unjamming alone does not ensure rupture—it is possible to have cells that are unable to invade a channel and rupture cell-cell adhesions (e.g., owing to low motility) but still have sufficient motility to fluidize the tissue. This includes, e.g., the points located in the lower center region of the (*ω*, Pe) plane. Therefore, we conclude that unjamming is necessary but not sufficient to create ruptures.

## V. DISCUSSION

To investigate the mechanism of how cells dissociate from collectively migrating strands in confinement, we re-analyzed data from our previous experiments [16], and used these results to motivate a phase-field simulation of invasion. We found in our experimental data that while larger channels led to larger cluster sizes, the most common outcome was always a single-cell dissociation event. To reproduce this effect within our model, we had to introduce the possibility that cells at the invasive front might spontaneously develop a leader state with a stronger-self-propulsion. Experimental measurements found no key difference between the time to rupture (survival time curves) for different channel sizes— opposite to the naive expectation that ruptures should be more common in channels with more cells. We were able to explain this feature of the data by assuming the cell adhesion weakens slightly in narrower channels relative to wider channels—a decrease of 30% in the narrowest channels. This reflects earlier results showing that contractility increases (leading to E-cadherin disengagement) and adhesions are more dynamic in confinement [16, 38, 39], but shows that these effects need not be quantitatively dramatic to explain relatively large changes in rupture rates. We also identify key control factors within our model, showing that increases in cell-cell adhesion strength suppress rupture (as seen experimentally [16]), that the probability of cells becoming leaders changes the distribution of cluster sizes, and that increasing cell-wall adhesion can both permit rupture by permitting cells to invade into narrow channels, as well as suppressing it, by preventing cells from detaching from the wall.

Earlier work has broadly argued that the degree of jamming and the ability of the tumor cells to rearrange may reflect whether cells from that tumor will metastasize [38, 53, 54]. We study, within our model, both the jamming transition and the rupture transition, and find that unjamming of the tissue is *necessary*, but not sufficient to predict rupture—there are many parameter sets where tissues are fluid but cancer cells are unable to dissociate. The details of this result may depend on assumptions of our model about cell-cell adhesion and motility. Researchers have employed various models to investigate the unjamming transition in tissues as a function of different control variables [8, 9, 21, 48]. For example, studies using the vertex model [48] and the Voronoi model [8] have explored how cell shape index and motility influence the jamming transition. These models make predictions about the influence of cell-cell adhesion—but adhesion enters only as an effective interfacial tension term [8, 55], changing the ability of cells to deform, but not immediately penalizing cell-cell separation or sliding of cell-cell junctions. Our focus on non-confluent tissues forces us to include cell-cell adhesion in a different form. We find that, in our phase field model, increasing cell-cell adhesion leads to less rearrangement and increased jamming. However, even within phase field models, there is no universal form of the cell-cell adhesion model, with Refs. [9, 20, 21, 28] all including adhesion in slightly different forms, and [7] disregarding it entirely, while [10] do not include a cell-cell adhesive force but do include the effect of a cell-cell friction, which also changes dynamics of rearrangement. Our model could also be extended to include more explicit dynamics of bond rupture [41, 56, 57]. Broadly, our results that increasing cell-cell adhesion leads to jamming is supported by experiments, but with some contradictory results. Many experiments support the idea that decreasing cell-cell adhesion can lead to tissue fluidization [14, 52, 58, 59] consistent with our observation. However, unjamming may also co-occur with the amplification of local adhesive intercellular stresses in human bronchial epithelial cells [48].

Our work shows that increasing cell-cell adhesion strength decreases the rate of rupture, but—if that rupture happens—increases the average size of the cluster that dissociates. This result may be relevant to the larger re-evaluation of the role of E-cadherin in cancer. The loss of intercellular adhesion molecule E-cadherin has long been considered a hallmark when tumor cells transition into an invasive phenotype resulting in cancer metastasis [60]. However, recent experiments have shown that E-cadherin expression is paradoxically correlated with cancer metastasis—while loss of E-cadherin increases invasion, it also reduces cancer cell proliferation and survival rates [60, 61]. Larger dissociation events may also be an aspect of this puzzle, because large-cluster dissociations may be more successful metastases [5].

We note that in our simulations at default parameters, all the survival probabilities drop to almost 0 [Fig. 5(b)], while the experimental survival curves saturate to approximately 30% [Fig. 2(b)]. This discrepancy in final survival probability might be attributed to many potential factors. First, the chemical gradient in the experiments of [16] is created by simple diffusion and will become more shallow over time, potentially decreasing the strength of chemotaxis or abolishing it entirely. Decreasing chemotactic strength leads to the flattening of survival curves in our model [Fig. 7(d)]. Second, this variation in outcome could also potentially reflect channel-to-channel variability in the experiment, e.g., in geometry or fibronectin coating. Variation in cell-wall or cell-cell adhesion from channel to channel could result in some channels that have much lower rates of cell rupture. Finally, cell-cell adhesion strength could be increasing over the experiment time [62].

Our model simply assumes the polarity of cells follows a random reorientation, biased by the chemoattractant gradient [Eq. (3)]. This neglects any possible co-ordination of cell polarities between neighboring cells, e.g., alignment of cell polarity to neighbors [63], or contact inhibition of locomotion (CIL) leading to repolarization [20, 29, 64], or many others reviewed in Ref. [65]. We view our assumptions here as a first minimal approach, but careful tests of cell-cell interaction may require us to extend our model. For instance, Ref. [33] shows that relatively rare populations of invasive leader cells can drive the invasion of non-leader cells, which could potentially be captured by this sort of cell-cell guidance [12, 66].

Our work has focused on the rupture and dissociation in collective cancer invasion, but it may have interesting connections to fracture events in other biological systems. Reproduction by fission in the metazoan *Trichoplax adherens* occurs by a motility-driven ductile-to-brittle transition of their epithelial layers [67], which might also reflect some of the control parameters we identify here. Fracture of epithelial monolayers under strain has been shown to depend on both keratin organization within the cell and cell-cell adhesive bond rupture [41]. Epithelial fracture has also been modeled within a vertex model with a detachment energy [42], predicting that the transition between the ability of the tissue to rearrange under strain and its tendency to break depends on the effective interfacial tension and contractility of the cells. Our model may also reflect this difference—but in our case, ruptures are not induced by a strain applied to the tissue but self-generated by the cells’ noisy motility. Formation of holes and voids can also occur in the spongy mesophyll, resulting in stable porous cell networks [68], or driven by cell contractility in epithelioid tissues [69]. An exciting future line of research is to understand to what extent and at what rate fracture events of all these various types can occur spontaneously, driven by randomness in cell motility and by chemical gradients, as observed in our work.

## ACKNOWLEDGMENTS

The authors acknowledge support from NIH Grants R35GM142847 (WW, EPI, and BAC) and R01GM142175 (RAL and KK). This work was carried out at the Advanced Research Computing at Hopkins (ARCH) core facility (rockfish.jhu.edu), which is supported by the National Science Foundation (NSF) Grant number OAC 1920103. We thank Yongtian Luo for a close reading of the manuscript.

## Appendix A: Alternate models

In this section, we present various models with different assumptions we have tested and show the process of how we reach our final model given in the main text.

### 1. Initial minimal model

As a starting point, we implement the simplest model with fixed intercellular adhesion *ω* = *ω*_0_ and only two cell states. We designate cells inside the channel or near the channel entrance as guided cells, while all other cells are classified as follower cells. The statistical results of our simulations are shown in Fig. 10(a) and 10(b). We can see that the average cluster sizes are larger than those observed in the experiments, and single-cell ruptures are not predominant in all channels. Additionally, the survival curves decline much more slowly compared to the experiments and do not collapse. By intuition, we would expect the survival probability to decrease with increasing channel width. Surprisingly, we notice that the 10 *μ*m curve is higher than the other curves. We will discuss this fact later in Appendix B.

### 2. Downregulating intercellular adhesion

In the experiments, we observe a good collapse of all survival curves for different channel widths on top of each other. However, the survival curves of our initial model in Fig. 10(b) do not collapse. We attempt to address this issue by reducing intercellular adhesion in narrow channels, as motivated by earlier experimental work [16]. With a reduction of only 30% in 6- and 10-micron channels and 20% in 20-micron channels, we achieve the collapse of all survival curves [see Fig. 10(d)], but the cluster-size distribution and the average cluster size [Fig. 10(c)] are still qualitatively different from the experimental data, with only rare single-cell ruptures.

### 3. Introducing leader cells

Compared to the initial model, here we introduce the high-metabolic-activity leader cells at the leading edge into the model, but do not tune cell-cell adhesion as a function of channel width. We notice that single-cell ruptures are predominant now [see Fig. 10(e)]. As for the survival probability, they decrease more rapidly with time, but still do not collapse onto a single curve [Fig. 10(f)]. Again, we see that the 10-micron channel has the longest survival curves, which we address in Appendix B.

### 4. Final model

Our final model featured in the main text combines the adjustments made in Models 2 and 3: downregulating intercellular adhesion in narrow channels and introducing leader cells, which recapitulates both the cluster-size distribution and the collapse of survival curves observed in experiments.

## Appendix B: How does cell size affect ruptures?

By intuition, we would expect the survival probability to decrease with increasing channel width. However, as we can see in Fig. 10(b) and 10(f), the 10-micron channel has slower ruptures than other channels. The reasoning behind the slower ruptures in the 10-micron channel lies in the proximity of its size to the default cell size (preferred radius *R* = 7 *μ*m) in our model. In the 10-micron channel, cells exhibit a broader cell-cell contact line compared to the 6-micron channel, making it harder to have ruptures—but in the 20-micron or wider channels, multiple cells can enter at once, so ruptures do not require breaking a larger contact line. Within our rough guiding hypothesis, we think of the 10-micron channel as having a higher initial energy barrier.

To verify our hypothesis, we first define the median survival time as the “half-life” of a survival curve *S*(*t*), i.e., the time when *S*(*t*) = 0.5, or more explicitly,

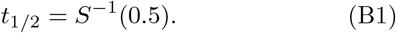

For the default *R* = 7 *μ*m cells, we observe a clear peak of *t*_1*/*2_ [extracted from Fig. 10(b)] at *W* = 10 *μ*m in Fig. 11(c). If our hypothesis holds true, the position of the peak should always be around the cell diameter 2*R*. Therefore, we next vary the preferred cell radius *R* to see how the peak of median survival time *t*_1*/*2_ changes. Figure 11(a) and 11(b) present the survival probabilities of the initial model for *R* = 3.5 *μ*m and *R* = 10 *μ*m, respectively. We notice that in Fig. 11(b), the 6-micron channels are too narrow for the cells with a diameter of ∼ 20 *μ*m to enter, resulting in the survival probability remaining at 1. After computing the corresponding median survival times, we find that the channel width where *t*_1*/*2_ peaks is indeed around the cell diameter and increases with cell size [Fig. 11(c)], which confirms our thoughts.

## Appendix C: Initial conditions

We first consider the static solution to the equation of motion [Eq. (1)] for an isolated cell in one dimension *ϕ*(*x*), so the active polarity vector **P** = 0, ℋ_exclusion_ = 0, and all adhesion terms vanish. If the area of the cell is fixed, then the area deviation term can also be ignored. Thus Eq. (1) degenerates to *δ*H_CH_*/δϕ* = 0 now, and the corresponding Euler-Lagrange equation of H_CH_ is [23]:

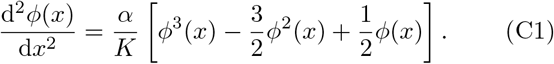

A domain wall can be introduced by forcing the two sides of the field to be in different phases, e.g., requiring *ϕ*(*x* →−∞) = 0 and *ϕ*(*x*→∞) = 1 as boundary conditions. By using the identity d^2^ tanh(*ax*)*/*d*x*^2^ = −2*a*^2^sech^2^(*ax*) tanh(*ax*), it can be readily verified that the profile:

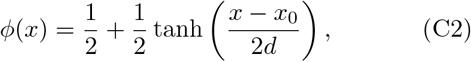

is a solution to the above nonlinear differential equation [Eq. (C1)] adhering to the boundary conditions, provided 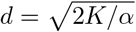 and *x*_0_ represent the width and position of the domain wall. Since our phase field model is in two dimensions, we use

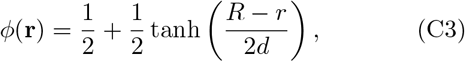

as the initial configuration, where *r* = |**r** −**r**_0_| is the distance to the center of the cell **r**_0_. The geometric-confinement field *ϕ*_wall_(**r**) is defined by a series of logistic sigmoid functions in a similar manner.

## Appendix D: Numerical details

By evaluating the functional derivative of the total Hamiltonian [Eq. (2)], the equation of motion [Eq. (1)] can be written as

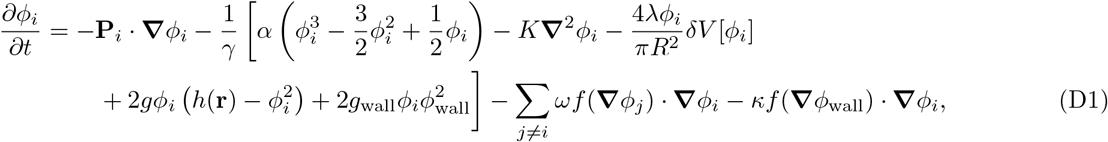

where we have introduced another auxiliary field 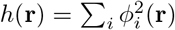 for parallel computing purpose (the simulation `C++` code is parallelized on multiprocessors by `Open MPI`) [7]. The area deviation term is 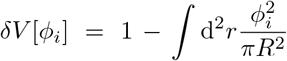, and we normalize the gradient via a saturation function 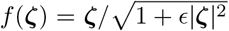 to prevent numerical instability resulting from very steep gradient [21]. Then, we employ forward differences to evolve the phase field equation for each cell, i.e.,

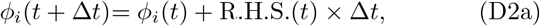

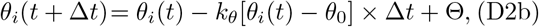

where R.H.S.(*t*) is the right-hand side of Eq. (D1), the noise Θ ∼ 𝒩 (0, 2*D*_*r*_Δ*t*), and the preferred direction *θ*_0_ points toward or along channels (*θ*_0_ = *π/*2). When computing the gradient ***∇****ϕ* and Laplacian ***∇***^2^*ϕ*, we apply first- and second-order central differences, respectively. We have matched all default parameter values in our model with the experiments, which are given in Table I.

During the simulation, we first implement a “thermalization” process for *T*_therm_ = 5 hours which allows all cells to evolve from the initial conditions we imposed and reach more natural shapes. After this process, we open the channels and keep on watching for *T*_tot_ = 18 hours, consistent with the experiments. Note that while we implement periodic boundary conditions in both the *x* and *y* directions, there is also a bottom wall, making the channels effectively closed at the top. We expect that the presence of this wall does not influence our core statistical results on dissociations. In our simulations, ruptures generally occur before cells hit the wall, and all channels generally have a rupture (Fig. 5)—except when motility is slowed (low Pe), in which case cells do not even reach the end of the channel. Once the first cells of the invading strand cells hit the wall, this suppresses dissociation—similar to if we had ended the simulation when cells impact the wall or removed them from the system, the other plausible boundary conditions. However, the presence of a wall (like the presence of a finite channel size in the experiments of [16]) could impact the very long-time dynamics in cases when dissociation is slow. Our simulated system size (channel length of 200 microns) is slightly smaller than the experimental channels (∼ 400 microns); this effect, like our finite number of cells, could in principle reduce the likelihood of extremely large clusters being found, though we have seen no evidence of this in the experiment.

The initial magnitudes of polarity for all cells are set to *p*_0_. Cells become guided once they get close enough to the channel. We define a box of (*W* + 24 *μ*m) × 6 *μ*m at the channel entrance in the simulation. Once the center of mass of a cell crosses into this region or inside the channel, the cell is considered a guided cell (*k*_*θ*_ ≠0), which can sense the chemical gradient and move with a larger velocity (magnitude of polarity set to 2*p*_0_). We also introduced leader cells at the invading front in our model—cells have a probability of becoming leader cells when they reach the front (become a tip cell). First, we have to define what a tip cell is: If there is any point of the cell surface where there is not a cell in the monolayer immediately above it, it is at the tip. In our model, whenever a guided cell becomes a tip cell, there is a probability for it to transition into a leader (magnitude of polarity set to 3*p*_0_). We give a sketch of the algorithm for this process in Algorithm 1. As discussed in the main text, the probability of becoming a leader cell inversely scales with its contact length with other cells in our model— cells are more likely to become leader cells if they have a lot of free space. Based on the form of the adhesion term in the model, we count a grid point at position **r** as a contact point of cell *i* when |***∇****ϕ*_*i*_(**r**) · _*j?*=*i*_ ***∇****ϕ*_*j*_(**r**)| = |***∇****ϕ*_*i*_ (***∇****h* −***∇****ϕ*_*i*_) | *>* 0.01 *μ*m^*−*2^. Therefore, Eq. (4) is calculated as *P* ^leader^ = max(1 − *n*_*i*_*/n*_*c*_, 0) in practice, where *n*_*i*_ is the number of contact points of cell *i*, and *n*_*c*_ is set to 256 in our simulation by default. Also, we note that in Algorithm 1 when a leader cell is no longer at the tip, it will revert to a guided cell state.

### Algorithm 1

How to identify cell states

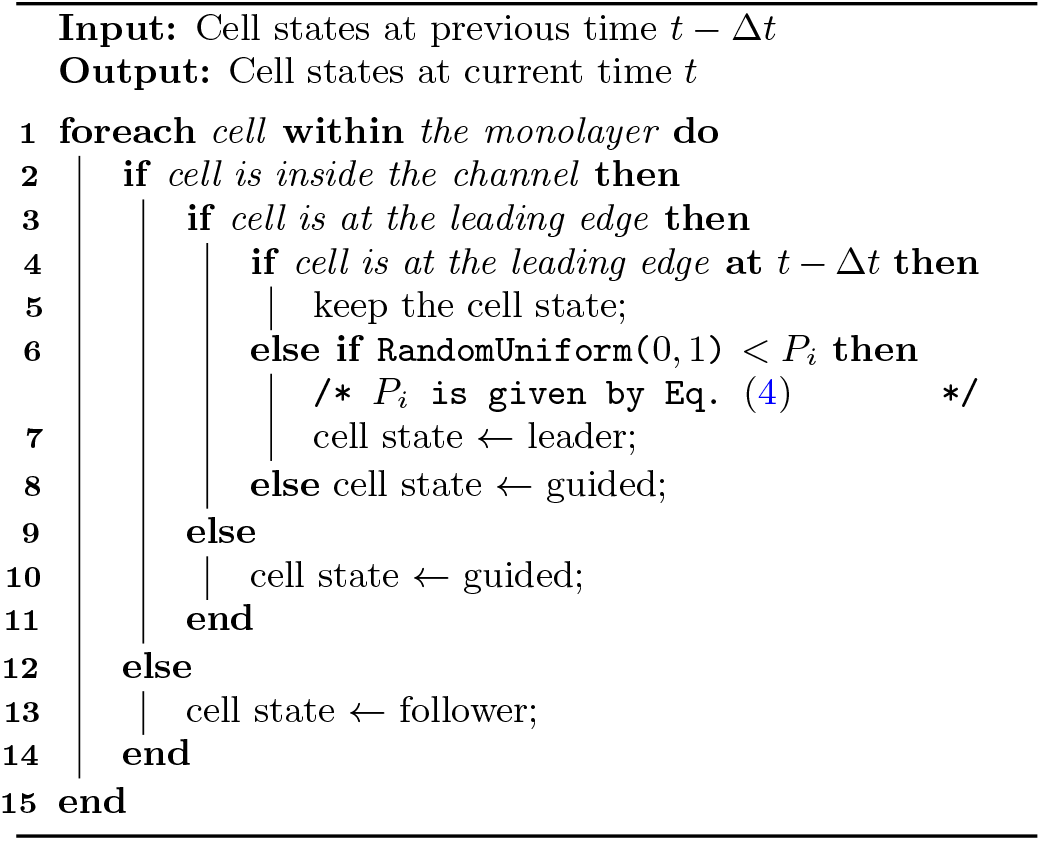

To improve computational efficiency, we also implement domain decomposition [7, 19, 20, 25], where we solve for each *ϕ*_*i*_ only within a smaller subdomain that shifts with the center of mass 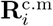. of each cell. The center of mass position for each cell is computed using the following equation:

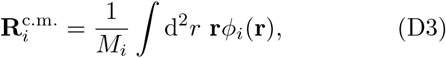

where *M*_*i*_ := ∫d^2^*r ϕ*_*i*_(**r**) is the total “mass” of the cell. Thus the center of mass velocity is readily given by 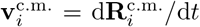. The mean squared displacement mentioned in Sec. IV G is computed by [7]

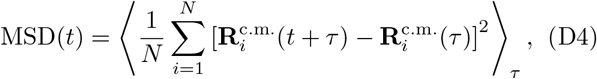

where the average ⟨· · · ⟩ is taken over time *τ* ∈ [0, *T*_tot_ −*t*]. We apply standard Kaplan-Meier survival analysis [18] to handle the data and generate survival probability plots using the `lifelines.KaplanMeierFitter` class provided by the Python survival-analysis package `lifelines` [70]. The 95% confidence intervals are computed using the exponential Greenwood confidence interval method [71].

## Appendix E: Supplemental simulation movies

### Movies 1–4

Dissociations of collectively invading monolayers in microchannels with different widths. Snapshots of these movies are presented in Fig. 4. All parameters are set to default values in these movies. Scale bar, 15 *μ*m.

### Movie 5

Cells can’t enter the narrow microchannel without cell-wall adhesion. Microchannel width *W* = 6 *μ*m. All parameters are set to default values except the cell-wall adhesion strength *κ* = 0. Scale bar, 15 *μ*m.

### Movies 6–7

Cells in jammed and unjammed states. Snapshots of these movies are presented in Fig. 8. Péclet numbers are Pe = 0.14 for the jammed state and 1.43 for the unjammed state. Scale bar, 15 *μ*m.

## Appendix F: Additional figures

Within the main paper (e.g., Fig. 2 and Fig. 5), we show the cluster size histograms with some cluster sizes grouped together, for legibility and clarity. In this section, we provide the full histograms. We also show the scatter plots of cluster size vs. rupture time for different channel widths.

**FIG. S1.**
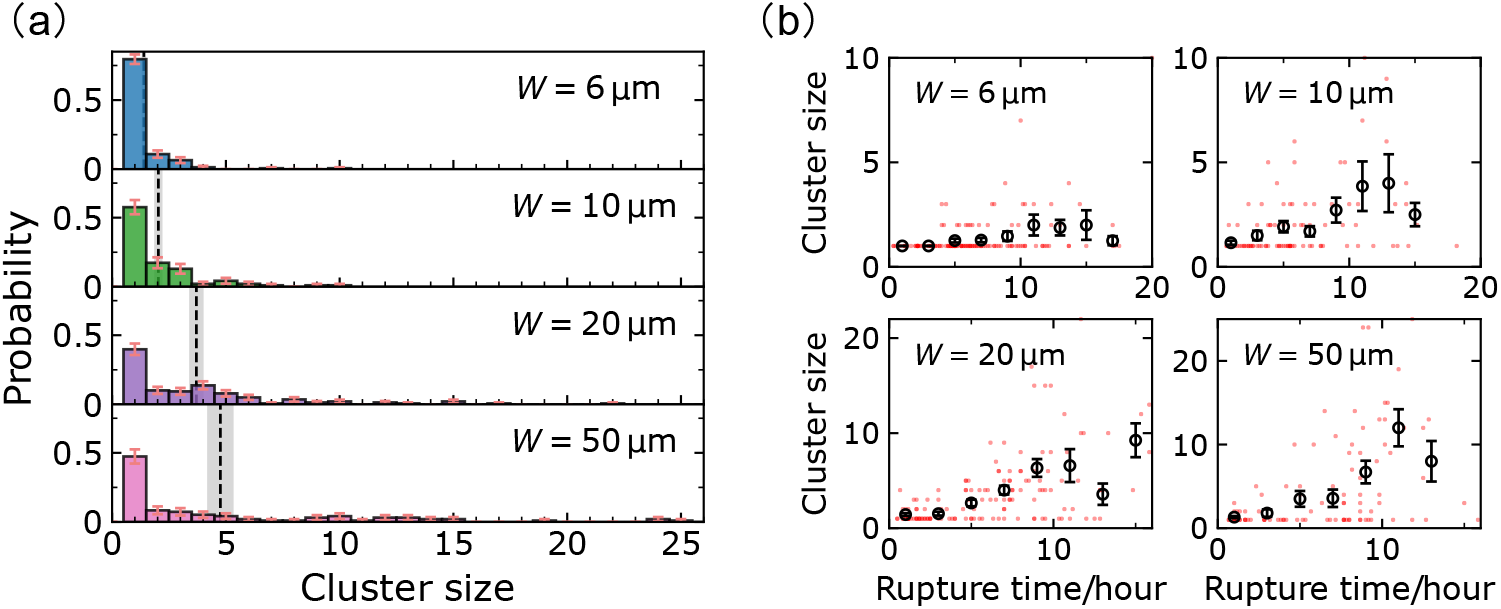
Experimental results. (a) Cluster-size distributions for microchannels of different widths. The dashed lines and gray areas denote the average cluster sizes *A* and their corresponding standard errors of the mean (SEM, or just SE). The error bars of the probabilities are determined by the binomial interval formula 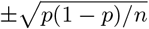. (b) Scatter plots exhibit the (rupture time, cluster size) phase plane. The circles represent the average cluster sizes within each time bin (2 hours), and the error bars indicate the corresponding SE. The standard error of the mean (SE) is calculated as 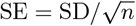, where SD is the standard deviation and *n* is the sample size.

**FIG. S2.**
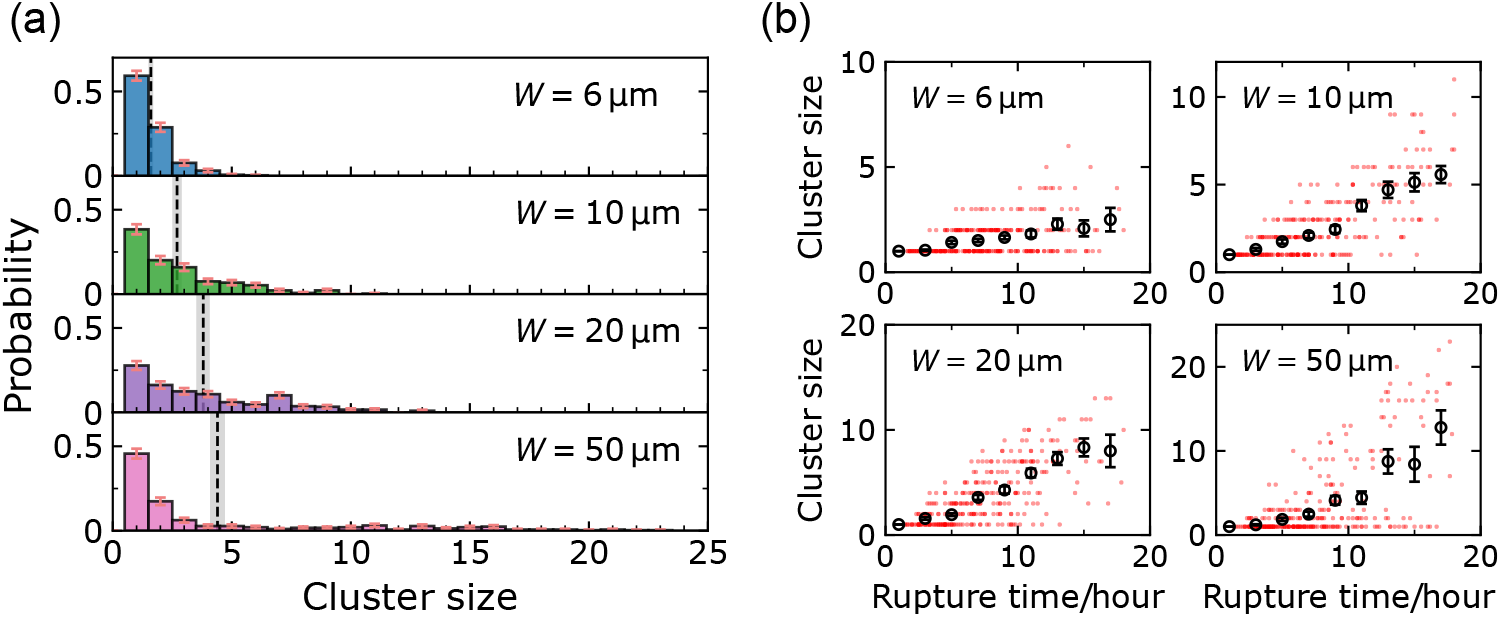
Simulation results with *ω* = 0.7*ω*_0_ for narrow channels, *ω* = 0.8*ω*_0_ for 20 *μ*m channels, and *ω* = *ω*_0_ for 50 *μ*m channels. (a) Cluster-size distributions for different channel widths. The dashed lines and gray areas denote the average cluster sizes *A* and their corresponding standard errors of the mean (SE). Panel (b) displays the (rupture time, cluster size) phase plane scatter plots. The circles represent the average cluster sizes within each time bin (2 hours), and the error bars indicate the corresponding SE.

**FIG. S3.**
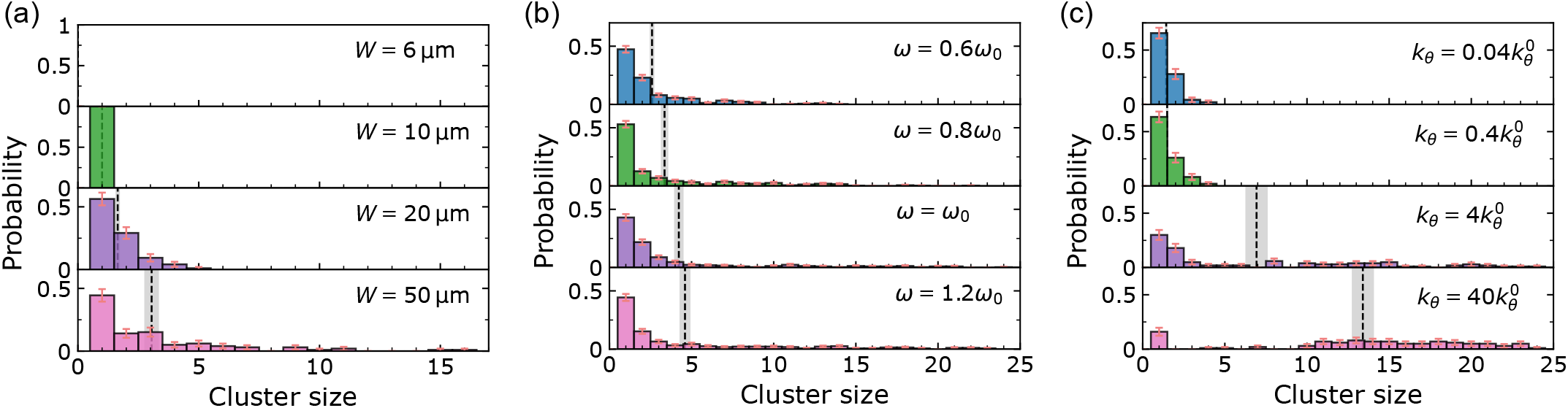
Simulation results. (a) Histograms when cell-wall adhesion *κ* is set to 0. (b) Histograms for different adhesion strength *ω* in 50 *μ*m channels. (c) Histograms for different chemotaxis strength *k*_*θ*_ in 50 *μ*m channels. The dashed lines and gray areas in (a)–(c) denote the average cluster sizes *A* and their corresponding SE.

## Notes

### Competing Interest Statement

The authors have declared no competing interest.

## References

[1] M. Poujade, E. Grasland-Mongrain, A. Hertzog, J. Jouanneau, P. Chavrier, B. Ladoux, A. Buguin, and P. Silberzan, Proceedings of the National Academy of Sciences 104, 15988 (2007).

[2] M. Chuai, D. Hughes, and C. J. Weijer, Current Genomics 13, 267 (2012).

[3] C. J. Weijer, Journal of cell science 122, 3215 (2009).

[4] A. Haeger, M. Krause, K. Wolf, and P. Friedl, Biochimica et Biophysica Acta (BBA) - General Subjects 1840, 2386 (2014).

[5] K. J. Cheung and A. J. Ewald, Science 352, 167 (2016).

[6] B. J. Green, M. Marazzini, B. Hershey, A. Fardin, Q. Li, Z. Wang, G. Giangreco, F. Pisati, et al., Small 18, 2106097 (2022).

[7] B. Loewe, M. Chiang, D. Marenduzzo, and M. C. Marchetti, Phys. Rev. Lett. 125, 038003 (2020).

[8] D. Bi, X. Yang, M. C. Marchetti, and M. L. Manning, Phys. Rev. X 6, 021011 (2016).

[9] G. Zhang, R. Mueller, A. Doostmohammadi, and J. M. Yeomans, Journal of The Royal Society Interface 17, 20200312 (2020).

[10] M. Chiang, A. Hopkins, B. Loewe, D. Marenduzzo, and M. C. Marchetti, arXiv preprint arXiv:2403.10715 10.48550/arXiv.2403.10715 (2024).

[11] A. Boromand, A. Signoriello, F. Ye, C. S. O’Hern, and M. D. Shattuck, Physical Review Letters 121, 248003 (2018).

[12] V. Tarle, E. Gauquelin, S. R. K. Vedula, J. D’Alessandro, C. T. Lim, B. Ladoux, and N. S. Gov, Phys. Biol. 14, 035001 (2017).

[13] E. M. Balzer, Z. Tong, C. D. Paul, W. C. Hung, K. M. Stroka, A. E. Boggs, S. S. Martin, and K. Konstantopoulos, The FASEB Journal 26, 4045 (2012).

[14] W.-J. Lin and A. Pathak, bioRxiv 10.1101/2023.04.10.536258 (2023).

[15] M. Saini, B. M. Szczerba, and N. Aceto, Cancer Research 79, 6067 (2019).

[16] R. A. Law, A. Kiepas, H. E. Desta, E. Perez Ipiña, M. Parlani, S. J. Lee, C. L. Yankaskas, R. Zhao, P. Mistriotis, N. Wang, Z. Gu, P. Kalab, P. Friedl, B. A. Camley, and K. Konstantopoulos, Science Advances 9, eabq6480 (2023).

[17] E. Perez Ipiña, J. d’Alessandro, B. Ladoux, and B. A. Camley, Proceedings of the National Academy of Sciences 121, e2318248121 (2024).

[18] E. L. Kaplan and P. Meier, Journal of the American statistical association 53, 457 (1958).

[19] M. Nonomura, PloS one 7, e33501 (2012).

[20] B. A. Camley, Y. Zhang, Y. Zhao, B. Li, E. Ben-Jacob, H. Levine, and W.-J. Rappel, Proceedings of the National Academy of Sciences 111, 14770 (2014).

[21] J. Löber, F. Ziebert, and I. S. Aranson, Scientific Reports 5, 1 (2015).

[22] B. Palmieri, Y. Bresler, D. Wirtz, and M. Grant, Scientific reports 5, 11745 (2015).

[23] R. Mueller, J. M. Yeomans, and A. Doostmohammadi, Phys. Rev. Lett. 122, 048004 (2019).

[24] A.-F. Bitbol and J.-B. Fournier, Phys. Rev. E 83, 061107 (2011).

[25] D. Shao, W.-J. Rappel, and H. Levine, Physical review letters 105, 108104 (2010).

[26] We note this force balance interpretation is not quite correct for our model, because the terms ∂ϕi/∂t adh are not derived from a Hamiltonian.

[27] F. L. H. Brown, Annu. Rev. Phys. Chem. 59, 685 (2008).

[28] P. Zadeh and B. A. Camley, Phys. Rev. E 106, 054413 (2022).

[29] D. A. Kulawiak, B. A. Camley, and W.-J. Rappel, PLOS Computational Biology 12, e1005239 (2016).

[30] P. Zadeh and B. A. Camley, arXiv preprint arXiv:2404.07390 (2024).

[31] N. Sepúlveda, L. Petitjean, O. Cochet, E. Grasland-Mongrain, P. Silberzan, and V. Hakim, PLOS Computational Biology 9, e1002944 (2013).

[32] E. Scarpa and R. Mayor, Journal of Cell Biology 212, 143 (2016).

[33] S. E. Leggett, M. C. Brennan, S. Martinez, J. Tien, and C. M. Nelson, Cellular and Molecular Bioengineering, 1 (2024).

[34] K. J. Cheung, E. Gabrielson, Z. Werb, and A. J. Ewald, Cell 155, 1639 (2013).

[35] P. Y. Hwang, A. Brenot, A. C. King, G. D. Longmore, and S. C. George, Cancer Research 79, 1899 (2019).

[36] J. Zhang, K. F. Goliwas, W. Wang, P. V. Taufalele, F. Bordeleau, and C. A. Reinhart-King, Proceedings of the National Academy of Sciences 116, 7867 (2019).

[37] S. SenGupta, C. A. Parent, and J. E. Bear, Nature Reviews Molecular Cell Biology 22, 529 (2021).

[38] O. Ilina, P. G. Gritsenko, S. Syga, J. Lippoldt, C. A. La Porta, O. Chepizhko, S. Grosser, M. Vullings, G.-J. Bakker, J. Starruß, et al., Nature cell biology 22, 1103 (2020).

[39] E. O. Wisniewski, P. Mistriotis, K. Bera, R. A. Law, J. Zhang, M. Nikolic, M. Weiger, M. Parlani, S. Tuntithavornwat, A. Afthinos, et al., Science advances 6, eaba6505 (2020).

[40] D. Bi, J. H. Lopez, J. M. Schwarz, and M. L. Manning, Soft Matter 10, 1885 (2014).

[41] J. Duque, A. Bonfanti, J. Fouchard, L. Baldauf, S. R. Azenha, E. Ferber, A. Harris, E. H. Barriga, A. J. Kabla, and G. Charras, bioRxiv, 2023 (2023).

[42] Y. Chen, Q. Gao, J. Li, F. Mao, R. Tang, and H. Jiang, Phys. Rev. Lett. 128, 018101 (2022).

[43] S. Gupta, A. E. Patteson, and J. M. Schwarz, New J. Phys. 23, 093042 (2021).

[44] N. G. Van Kampen, Stochastic processes in physics and chemistry, Vol. 1 (Elsevier, 1992).

[45] B. A. Camley, Journal of Physics: Condensed Matter 30, 223001 (2018).

[46] B. A. Camley and W.-J. Rappel, Physical Review E 89, 062705 (2014).

[47] J. A. Mitchel, A. Das, M. J. O’Sullivan, I. T. Stancil, S. J. DeCamp, S. Koehler, J. P. Butler, J. J. Fredberg, M. A. Nieto, D. Bi, and J.-A. Park, bioRxiv 10.1101/665018 (2019).

[48] J.-A. Park, J. H. Kim, D. Bi, J. A. Mitchel, N. T. Qazvini, K. Tantisira, C. Y. Park, M. McGill, S.-H. Kim, B. Gweon, et al., Nature materials 14, 1040 (2015).

[49] C. Malinverno, S. Corallino, F. Giavazzi, M. Bergert, Q. Li, M. Leoni, A. Disanza, E. Frittoli, A. Oldani, E. Martini, et al., Nature materials 16, 587 (2017).

[50] E. Lawson-Keister and M. L. Manning, Current Opinion in Cell Biology 72, 146 (2021).

[51] A. Mongera, P. Rowghanian, H. J. Gustafson, E. Shelton, D. A. Kealhofer, E. K. Carn, F. Serwane, A. A. Lucio, J. Giammona, and O. Campàs, Nature 561, 401 (2018).

[52] E. Blauth, H. Kubitschke, P. Gottheil, S. Grosser, and J. A. Käs, Frontiers in Physics 9, 666709 (2021).

[53] S. Grosser, J. Lippoldt, L. Oswald, M. Merkel, D. M. Sussman, F. Renner, P. Gottheil, E. W. Morawetz, T. Fuhs, X. Xie, et al., Physical Review X 11, 011033 (2021).

[54] P. Gottheil, J. Lippoldt, S. Grosser, F. Renner, M. Saibah, D. Tschodu, A.-K. Poßögel, A.-S. Wegscheider, B. Ulm, K. Friedrichs, et al., Physical Review X 13, 031003 (2023).

[55] M. L. Manning, R. A. Foty, M. S. Steinberg, and E.-M. Schoetz, Proceedings of the National Academy of Sciences 107, 12517 (2010).

[56] S. Weng, R. J. Huebner, and J. B. Wallingford, Cell Reports 39 (2022).

[57] E. McEvoy, T. Sneh, E. Moeendarbary, Y. Javanmardi, N. Efimova, C. Yang, G. E. Marino-Bravante, X. Chen, J. Escribano, F. Spill, et al., Nature communications 13, 7089 (2022).

[58] M. Sadati, N. T. Qazvini, R. Krishnan, C. Y. Park, and J. J. Fredberg, Differentiation 86, 121 (2013).

[59] D. T. Tambe, C. Corey Hardin, T. E. Angelini, K. Rajendran, C. Y. Park, X. Serra-Picamal, E. H. Zhou, M. H. Zaman, J. P. Butler, D. A. Weitz, et al., Nature materials 10, 469 (2011).

[60] G. C. Russo, A. J. Crawford, D. Clark, J. Cui, R. Carney, M. N. Karl, B. Su, B. Starich, T.-S. Lih, P. Kamat, et al., Oncogene, 1 (2024).

[61] V. Padmanaban, I. Krol, Y. Suhail, B. M. Szczerba, N. Aceto, J. S. Bader, and A. J. Ewald, Nature 573, 439 (2019).

[62] S. Garcia, E. Hannezo, J. Elgeti, J.-F. Joanny, P. Silberzan, and N. S. Gov, Proceedings of the National Academy of Sciences 112, 15314 (2015).

[63] T. Vicsek, A. Cziróók, E. Ben-Jacob, I. Cohen, and O. Shochet, Phys. Rev. Lett. 75, 1226 (1995).

[64] W. Wang and B. A. Camley, Phys. Rev. E 109, 054408 (2024).

[65] B. A. Camley and W.-J. Rappel, Journal of physics D: Applied physics 50, 113002 (2017).

[66] A. J. Kabla, Journal of The Royal Society Interface 9, 3268 (2012).

[67] V. N. Prakash, M. S. Bull, and M. Prakash, Nature Physics 17, 504 (2021).

[68] J. D. Treado, A. B. Roddy, G. Théroux-Rancourt, L. Zhang, C. Ambrose, C. R. Brodersen, M. D. Shattuck, and C. S. O’Hern, Journal of the Royal Society Interface 19, 20220602 (2022).

[69] J.-Q. Lv, P.-C. Chen, Y.-P. Chen, H.-Y. Liu, S.-D. Wang, J. Bai, C.-L. Lv, Y. Li, Y. Shao, X.-Q. Feng, et al., Nature Physics, 1 (2024).

[70] C. Davidson-Pilon, Journal of Open Source Software 4, 1317 (2019).

[71] M. Greenwood, Reports on public health and medical subjects 33, 1 (1926).

